# Rejection of inappropriate synaptic partners mediated by transcellular FLRT2-UNC5 signaling

**DOI:** 10.1101/2022.08.29.505771

**Authors:** Cameron L. Prigge, Arsha Sharma, Mayur Dembla, Malak El-Quessny, Christopher Kozlowski, Caitlin E. Paisley, Tyler Johnson, Luca Della Santina, Marla B. Feller, Jeremy N. Kay

## Abstract

During nervous system development, neurons choose synaptic partners with remarkable specificity; however, the cell-cell recognition mechanisms governing rejection of inappropriate partners remain enigmatic. Here we show that mouse retinal neurons avoid inappropriate partners using the FLRT2-UNC5 receptor-ligand system. Within the inner plexiform layer (IPL), FLRT2 is expressed by direction-selective (DS) circuit neurons, whereas UNC5C/D are expressed by non-DS neurons projecting to adjacent IPL sublayers. In vivo gain- and loss-of-function experiments demonstrate that FLRT2-UNC5 binding eliminates growing DS dendrites that have strayed from the DS circuit IPL sublayers. Abrogation of FLRT2-UNC5 binding allows mistargeted arbors to persist, elaborate, and acquire synapses from inappropriate partners. Conversely, UNC5C misexpression within DS circuit sublayers inhibits dendrite growth and drives arbors into adjacent sublayers. Mechanistically, UNC5s promote dendrite elimination by interfering with FLRT2-mediated adhesion. Based on their broad expression, FLRT-UNC5 recognition is poised to exert widespread effects upon synaptic partner choices across the nervous system.

## INTRODUCTION

Our ability to perceive the world around us results from the exquisitely precise synaptic connections made between neurons during development. These connections establish neural circuits dedicated to specific computations and functions. During circuit development, growing neurons actively choose which cells to connect with and which to avoid using cell-surface receptors. Defects in this process can compromise circuit function, particularly when such defects produce connections between the wrong partners (Sanes and Zipursky, 2020). Despite their importance, the molecular mechanisms underlying synaptic partner choices remain enigmatic.

Molecular studies of synaptic specificity have mainly focused on positive cues, such as attractive or adhesive recognition molecules, that affirm appropriate connections. By contrast, far less is known about negative cues, which discourage inappropriate connections (Lefebvre et al., 2015; Prigge and Kay, 2018; Sanes and Zipursky, 2020). It is generally presumed that both positive and negative cues are important for selective cell-cell recognition during circuit formation. However, few circuit-specific negative cues that prevent cross-circuit wiring are currently known (Pecho-Vrieseling et al., 2009). Instead, most negative cues identified so far are repulsive guidance molecules that prevent axons and dendrites from growing into particular regions (Cang and Feldheim, 2013; Kolodkin and Tessier-Lavigne, 2011; Lefebvre et al., 2015; Pederick et al., 2021; Sanes and Zipursky, 2020). In many cases, deletion of such molecules can alter neurite targeting without disturbing the correct pairing of synaptic partners (Matsuoka et al., 2011; Sun et al., 2013). Therefore, it remains unclear whether negative cues are actually involved in partner selection, or if they are they are instead mainly involved in neurite guidance.

The retina is a useful model system to investigate synaptic partner choice due to its stereotyped laminar organization of synaptic connections. During development, individual retinal ganglion cells (RGCs) and their presynaptic partners, the bipolar and amacrine cells, project their axons or dendrites into just one or a few sublayers within the neuropil known as the inner plexiform layer (IPL; Fig. 1A). This laminar organization facilitates the cellular rendezvous necessary for circuit-specific connections to form. Here, to investigate the role of negative cues in establishing laminar and synaptic specificity, we used the mouse retinal direction-selective (DS) circuit (Fig. 1A) as our model system. This evolutionarily conserved circuit encodes the direction of image motion, thereby driving multiple visual behaviors that depend on motion detection (Fredericks et al., 1988; Giolli et al., 2006; Hamilton et al., 2021; Kim et al., 2022; Mauss et al., 2017; Patterson et al., 2022; Ray and Kay, 2015). Motion direction is encoded via the firing responses of direction-selective ganglion cells (DSGCs) that preferentially respond to stimuli moving in particular directions. These DS responses are generated via synaptic inputs onto DSGCs from specific types of bipolar and amacrine cells – especially the starburst amacrine cells, which provide extensive inhibitory inputs that are a particularly crucial determinant of the DS computation (Briggman et al., 2011; Fried et al., 2002; Hanson et al., 2019; Mauss et al., 2017; Yoshida et al., 2001).

**Figure 1:**
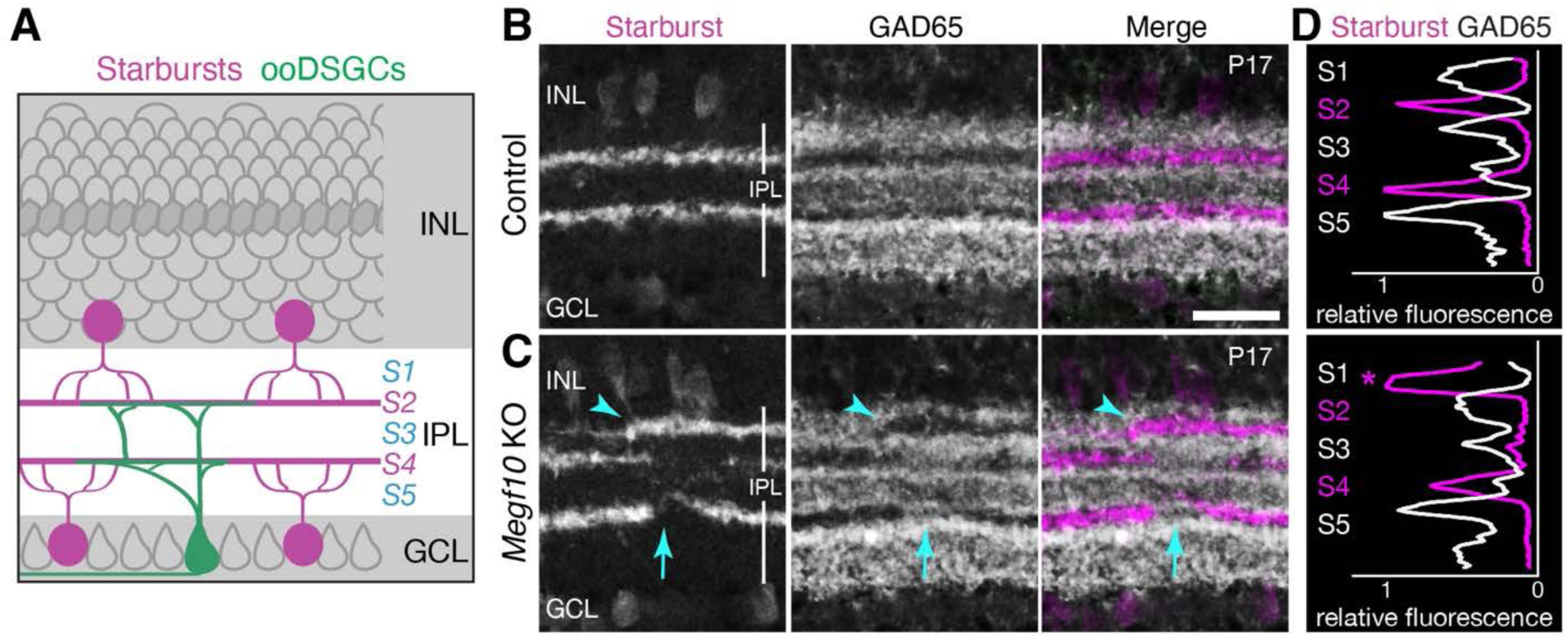
Mutual exclusion between DS circuit dendrites and neighboring GAD65^+^ dendrites. **A**. Schematic of mouse retina in cross-section view showing DS circuit neurons and their projections within inner plexiform layer (IPL) neuropil. Starburst amacrine cell bodies (magenta) are located in inner nuclear layer (INL; OFF cells) and ganglion cell layer (GCL; ON cells). OFF starbursts project to sublamina (S)2; ON starbursts project to S4. ON-OFF direction-selective ganglion cells (ooDSGCs, green) project bistratified dendritic trees to cofasciculate with starburst dendrites within S2 and S4. Starburst cells co-release both acetylcholine and GABA onto DSGCs. **B,C**. Retinal cross sections stained with anti-ChAT (starburst marker) and anti-GAD65, showing IPL phenotype in *Megf10* knockout (KO) animals. B, littermate controls (*Megf10^flox/flox^*; no cre). C, *Megf10* mutants (*Six3-Cre; Megf10^flox/flox^*). Controls (B) exhibit mutual exclusion between IPL territories expressing each marker. In mutants (C), starburst cells project inappropriately into IPL strata that are normally GAD65^+^, displacing GAD65 arbors (arrowheads). Starburst dendrites also sporadically fail to innervate regions within S2 and S4; GAD65+ arbors invade these starburst-free regions (arrows). Scale bar, 10 µm. **D**. Representative profile plots showing ChAT (magenta) and GAD65 (white) fluorescence across the IPL. Sublaminae S1-S5 are indicated. In control (top), peaks in GAD65 fluorescence align with troughs in ChAT fluorescence, and vice versa. Mutant trace shows starburst dendrites mistargeted to S1 (asterisk) and locally absent from S2. Nevertheless, peaks and troughs of each fluorescent signal are still aligned showing mutual exclusion is preserved.

The DS circuit occupies two dedicated IPL sublayers, denoted S2 and S4 (Fig. 1A), where DSGCs and starburst amacrine cells intertwine their dendritic arbors (amacrine dendrites have the unusual property of both sending and receiving synapses, so to distinguish them from conventional dendrites we will mainly refer to them as “arbors”). S2 encodes light OFF responses while S4 encodes ON responses. These IPL sublayers are assembled during development by affirmative cues that bring DS circuit partners together. First, adhesion and homotypic recognition mediate the stratification of starburst arbors, which occurs around postnatal day (P) 0 in mice (Ray et al., 2018; Stacy and Wong, 2003). Then, over the next few days, adhesion molecules serve to recruit ON-OFF (oo)DSGC dendrites and bipolar cell axons to the starburst laminar scaffold (Duan et al., 2014; Duan et al., 2018; Peng et al., 2017). Strict ooDSGC stratification within these sublayers ensures that the vast majority of inhibitory inputs onto ooDSGC dendrites are GABAergic synapses supplied by starburst amacrine cells (Bleckert et al., 2018; Pei et al., 2015; Sivyer et al., 2019; Vaney and Pow, 2000). Numerous other amacrine cell types project to IPL sublayers directly adjacent to the DS circuit, mere microns away, and would therefore be easily accessible to ooDSGC dendrites during synaptogenesis. Nevertheless, ooDSGCs avoid such connections. The mechanisms that prevent ooDSGCs from making synapses in adjacent non-DS layers remain unknown..

In this study we identify negative molecular cues that prevent ooDSGCs and starburst cells (which we collectively term “DS circuit neurons”) from forming connections in the wrong IPL sublayers. Fibronectin leucine-rich repeat transmembrane (FLRT) proteins are multifunctional cell-surface molecules that participate in cell adhesion, repulsion, and synaptogenesis (Klein and Pasterkamp, 2021). These effects are mediated by binding to multiple ligands of the latrophilin (LPHN) and Uncoordinated-5 (UNC5) protein families (Fig. 2A, C) (del Toro et al., 2020; Jackson et al., 2015; Jackson et al., 2016; Lu et al., 2015; O’Sullivan et al., 2012; Ranaivoson et al., 2015; Sando et al., 2019; Seiradake et al., 2014; Visser et al., 2015; Yamagishi et al., 2011). Adhesive and synaptogenic FLRT functions are mediated by LPHN binding; by contrast, the impact of binding to UNC5s remains uncertain. While FLRTs are known to be repulsive ligands for UNC5 receptors, it is unknown whether “reverse” UNC5-to-FLRT signaling can guide FLRT^+^ neurons or whether UNC5 binding influences FLRT-LPHN signaling in vivo. To investigate the function of the FLRT-UNC5 component of this tripartite molecular system, we took advantage of the developing IPL where we previously showed that FLRT2 and UNC5C are expressed in complementary patterns: FLRT2 levels are high in the DS circuit sublayers while UNC5C fills surrounding strata (Visser et al., 2015) (Fig. 2B,D). In stripe assays using cultured primary retinal neurons, UNC5C protein inhibits arbor growth by FLRT2^+^ cells (Visser et al., 2015). These results led us to hypothesize that FLRT2-UNC5C signaling could impact DS circuit laminar specificity in vivo.

**Figure 2:**
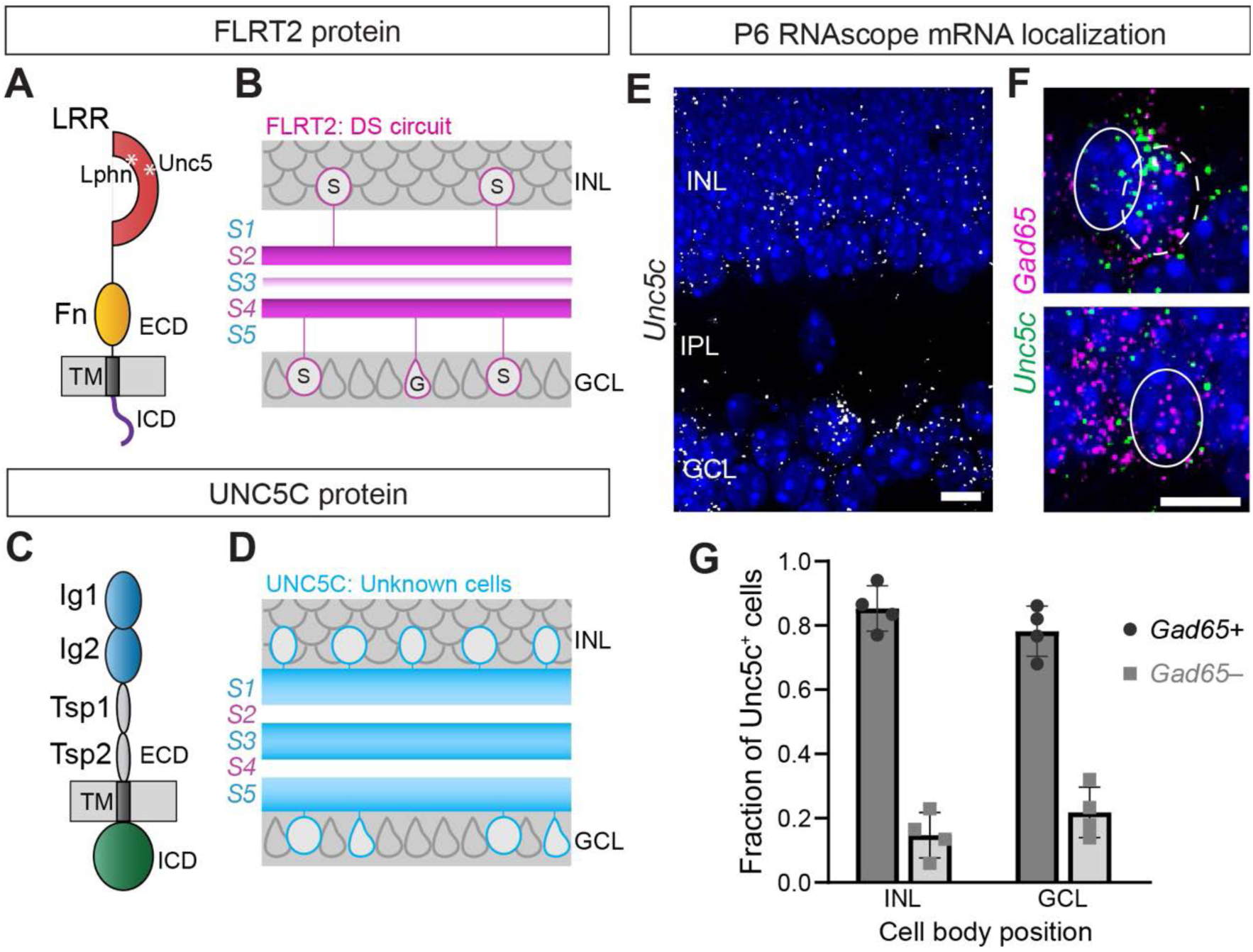
FLRT2 and UNC5C are poised to mediate mutual exclusion. **A,C**. Schematic of FLRT2 (A) and UNC5C (C) protein structure. FLRT2 has an extracellular leucine rich repeat (LRR) and Fibronectin (Fn)-like domain. Binding sites for latrophilins (Lphn) and UNC5s are indicated (asterisks). UNC5C has two immunoglobulin (Ig) and thrombospondin (Tsp) domains. TM, transmembrane domain; ECD, extracellular domain; ICD, intracellular domain. **B,D.** Summary of FLRT2 (B) and UNC5C (D) protein expression within the IPL, as shown in our previous study (Visser et al., 2015). DS circuit neurons including DS ganglion cells (G) and starburst cells (S) contribute prominently to FLRT2 expression pattern. Some protein is also localized to S3. UNC5C protein is abundant within S1, S3, and S5 but is absent from S2 and S4. **E-F**. RNAscope in situ hybridization using probes against *Unc5c* (E,F) and Gad65 (F). Blue, Hoechst nuclear counterstain. E: Overview of *Unc5c* expression showing prominent labeling of INL and GCL. F, top: Representative examples of INL cells expressing *Unc5c*. Some cells co-expressed Gad65 (dashed line) whereas others were Gad65-negative (solid line). F, bottom: Representative example of Gad65*+* cell that did not express *Unc5c* (solid line). Scale bars, 10 µm. **G.** Summary data showing fraction of *Unc5c^+^* cells co-expressing Gad65. Most *Unc5^+^* cells in GCL are GABAergic amacrines rather than RGCs. N = 4 animals.

Here we have built and deployed a wide range of mouse genetic tools, including a new mouse strain in which FLRT2-UNC5 binding is selectively disrupted, to demonstrate that the FLRT2-UNC5 receptor-ligand system plays a central role in DS circuit synaptic specificity. The study has two parts. First, we describe laminar and synapse specificity errors in mouse mutants lacking FLRT2 or UNC5-family molecules. Second, we determine the developmental origins and molecular mechanisms underlying these IPL targeting phenotypes. Unexpectedly, we identify in wild-type mice a brief developmental period when arbors of DS circuit neurons stray from the starburst scaffold. We find that FLRT2 and UNC5 are dispensable for initial IPL targeting, but that FLRT2-UNC5 interactions act as an “error correction” mechanism to constrain growth of these stray arbors. Further, we show that UNC5 binding constrains growth of FLRT2^+^ arbors by interfering with FLRT2-LPHN adhesion. Altogether, our results support a model in which affirmative mechanisms control the initial pairing of ooDSGC and starburst dendrites, whereas negative mechanisms – mediated by FLRT2-UNC5 binding – are needed during subsequent dendrite growth to prevent ooDSGC arbors from abandoning their synaptic partners.

## RESULTS

### Repulsion between adjacent circuits establishes DS circuit-specific IPL territories

To determine whether repulsion is involved in laminar specificity of DS circuit IPL projections, we used GAD65 immunolabeling to mark adjacent IPL sublayers. Antibodies to GAD65 label a diverse population of medium- and wide-field GABAergic amacrine cell types, most of which are narrowly stratified in individual IPL sublayers (Diamond, 2017; Pourcho and Goebel, 1983). Together, GAD65-immunoreactive amacrine arbors fill most of the IPL, with the exception of the DS circuit IPL sublayers marked by antibodies to the starburst markers vesicular acetylcholine transporter (VAChT) or choline acetyltransferase (ChAT; Fig. 1B). In addition to acetylcholine, starburst cells also release GABA; however, they express only the GAD67 form of glutamate decarboxylase (Famiglietti and Sundquist, 2010) (Supplementary Fig. 2S1). Therefore, GAD65 selectively marks GABAergic amacrine cells that are not part of the DS circuit.

To test the role of repulsion in establishing the mutually exclusive projection pattern of GAD65^+^ and DS circuit arbors, we utilized *Megf10* mutant mice to selectively alter DS circuit IPL projections (Ray et al., 2018). MEGF10 is a starburst-specific cell surface molecule; loss of this molecule causes focal disruptions in laminar targeting involving both gaps in starburst IPL innervation and displacement of starburst arbors into inappropriate sublayers (Fig. 1C). These starburst errors are precisely mirrored by ooDSGC dendrites, which strictly follow starburst arbors even when they are mistargeted (Ray et al., 2018). If repulsion exists between DS circuit and GAD65^+^ arbors, the two populations would be expected to occupy mutually exclusive territories even when DS circuit projections are altered. Consistent with this prediction, we found that GAD65^+^ arbors in *Megf10* mutants did not commingle with mistargeted starburst arbors that encroached on their territory. Instead, the presence of mistargeted starburst arbors displaced GAD65^+^ arbors from their usual IPL strata (Fig. 1B,C). Furthermore, gaps in the DS circuit IPL sublayers were inappropriately innervated by GAD65^+^ arbors, suggesting a defect in repulsion when starburst and/or ooDSGC arbors are absent (Fig. 1C). These results support the notion that repulsive cues prevent adjacent IPL sublayers from intermingling.

### FLRT2 and UNC5C are poised to mediate circuit segregation

We next sought to identify molecular cues that could mediate repulsion between the DS circuit and GAD65^+^ amacrine cells within the IPL. We focused on FLRT2 and its UNC5-family ligands because previous work established that FLRT2 mRNA and protein are abundantly expressed by starburst and ooDSGC arbors in sublayers S2 and S4 (Visser et al., 2015; Fig. 2B). In contrast, UNC5C, a FLRT ligand, localizes to non-DS IPL regions in a pattern resembling GAD65 immunoreactivity (Visser et al., 2015; Fig. 2D). To determine whether UNC5C is expressed by the GABAergic amacrine cells that project to IPL sublayers bordering the DS circuit we performed *in situ* hybridization for *Unc5c* and Gad65 (encoded by the *Gad2* gene). This analysis revealed that *Unc5c* is mainly expressed by Gad65*^+^* GABAergic amacrine cells (Fig. 2E-G; Supplementary Fig. 2S1). A subset of RGCs also expressed *Unc5c*, but these accounted for a small fraction of the *Unc5c^+^* neuronal population (Fig. 2G).

These data are consistent with bulk and single-cell transcriptomic datasets, in which we found that *Unc5c* was not expressed at appreciable levels by DS circuit neurons (Kay et al., 2012; Shekhar et al., 2022; Tran et al., 2019; Yan et al., 2020). Using single-cell data (Yan et al., 2020), we identified 24 *Unc5c-*expressing amacrine cell types, 23 of which were GABAergic (Supplementary Fig. 2S1B). Seven of these *Unc5c^+^*clusters correspond to well-characterized amacrine cell types with known projection patterns in the IPL; all seven project to IPL strata located adjacent to the DS circuit sublayers (Supplementary Fig. 2S1). Together, these results demonstrate that FLRT2 and UNC5C are expressed in a complementary manner by distinct cell types – i.e. DS circuit neurons and non-DS amacrine cells – with mutually exclusive stratification patterns in the IPL. Therefore, this receptor-ligand pair is positioned to provide repulsive laminar targeting cues during DS circuit development.

### DS circuit laminar targeting errors in *Flrt2* mutant mice

To test whether FLRT2-UNC5C signaling is required for DS circuit laminar targeting, we used a variety of approaches to disrupt this interaction and then assessed whether DSGC dendritic arbors stray from their normal S2 and S4 sublayers. First, we generated retina-specific *Flrt2* mutants (denoted *Flrt2^Ret^*), in which the *Six3-Cre* transgenic line (Furuta et al., 2000) was used to drive retinal recombination of a conditional *Flrt2^flox^* allele. Antibody labeling verified that FLRT2 protein was absent from the retinas of *Flrt2^Ret^* mice (Supplementary Fig. 3S1). To label ooDSGCs we used the *Hb9-GFP* transgene, which selectively marks ooDSGCs (Supplementary Fig. 3S2) that prefer ventral motion (Trenholm et al., 2011).

Staining for starburst and Hb9-DSGC arbors at P15-17 revealed two types of laminar targeting errors in *Flrt2^Ret^* animals, each of which occurred sporadically across mutant retina. First, a subset of Hb9-DSGC dendrites were uncoupled from the starburst scaffold, stratifying within ectopic IPL regions that did not contain starburst arbors (Fig. 3C-E). These laminar errors were highly specific: When not associated with the starburst scaffold, Hb9-DSGC dendrites instead targeted stereotyped ectopic locations in S1 and S3 (Fig. 3C,D,G,H). Some of these ectopic projections extended over large distances (Fig. 3H). We observed a similar phenotype in *Six3-Cre; Flrt2^flox/+^* heterozygotes, albeit at lower frequency (Supplementary Fig. 3S3). Ectopic arbors did not co-label with the axon marker Neurofilament-M, confirming their dendritic identity (Supplementary Fig. 3S4). Single-cell reconstructions revealed that ∼25% of *Flrt2* mutant Hb9-DSGCs made dendritic laminar errors (Fig. 3F,G, n = 3/12 cells projected ectopically). By contrast, *Flrt2^WT^* control animals rarely exhibited Hb9-DSGC dendrites outside the sublayers defined by starburst arbors (Fig. 3A,E).

**Figure 3:**
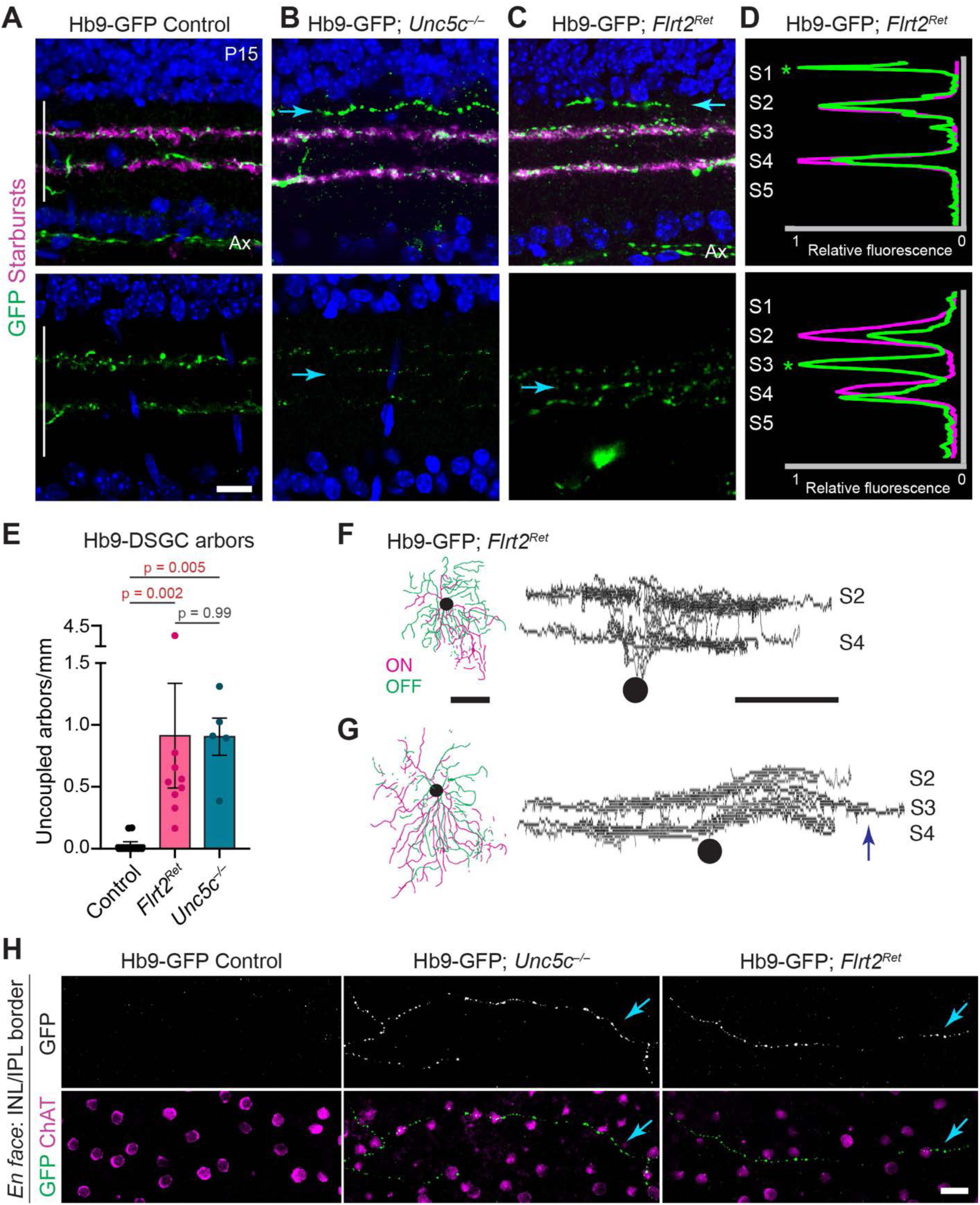
*Flrt2* and *Unc5c* are required for laminar targeting of ooDSGC dendrites. **A-C**. Representative retinal cross-sections from P15 Hb9-GFP mice of the specified *Unc5c* or *Flrt2* genotype, stained with anti-GFP to reveal ooDSGC arbors. Anti-Vacht staining (top row) labels starburst arbors. Blue, Hoechst nuclear counterstain. IPL (vertical bar) contains starburst and ooDSGC dendrites. In some sections, GFP^+^ axons (Ax) are also visible within nerve fiber layer. In littermate control mice carrying wildtype alleles of *Flrt2* and *Unc5c* (A), starburst and ooDSGC dendrites co-stratify within S2 and S4. In *Unc5c* null mutants (B) as well as *Flrt2^Ret^* mutants (C), ooDSGCs become uncoupled from the DS circuit scaffold, making laminar targeting errors into S1 (top row) and S3 (bottom row). Blue arrows indicate mistargeted dendrites. **D**. Representative fluorescence profile plots through the IPL, highlighting location of ectopic arbors (asterisks) in *Flrt2^Ret^* mutants. Also see Supplementary Fig. 3S6 for *Unc5c^−/−^* profile plots. **E**. Summary of ooDSGC laminar targeting errors, defined as number of GFP^+^ arbors that were uncoupled from starburst arbors and resided in ectopic sublayers. Control group pools wild-type littermates from *Flrt2* and *Unc5c* strains, as wild-type animals from each strain did not show phenotypic differences. Strength of ooDSGC phenotype was similar in *Flrt2^Ret^* and *Unc5c^−/−^* mutants. Statistics, Kruskal-Wallis test (main effect p = 9 x 10^-5^) followed by Dunn’s post-hoc test. P-values (shown on graph) were corrected for multiple comparisons. Red text denotes significant difference. Sample sizes (number of animals): littermate controls n = 11 (n = 6 *Flrt2^WT^;* n = 5 *Unc5c^+/+^*), *Flrt2^Ret^* n = 9, *Unc5c^−/−^* n = 5. Error bars, S.E.M. **F,G.** Single cell reconstructions of dye-filled ooDSGCs from *Hb9-GFP; Flrt2^Ret^* animals, showing an ooDSGC with normal S2-S4 stratification (F) and another cell projecting an ectopic arbor (arrow) into S3 (G). N = 3/12 filled cells exhibited ectopic arbors. Scale bar, 100 µm. **H**. Representative en face confocal images showing S1 ectopic ooDSGC arbors in *Unc5c^−/−^* and *Flrt2^Ret^* mutants. Images were acquired from retinal wholemounts in the plane of the IPL/INL border. Hb9-GFP^+^ arbors do not normally project to this location in wild-type retina (left) but are readily observed in both mutants (center, right; arrows). Scale bar, 20 µm. Scale bars, 20 µm (A-C, H); 100 µm (F, G).

To test whether this laminar targeting phenotype generalizes to additional ooDSGC subtypes, we utilized the *Drd4-GFP* transgenic line, which marks nasal motion-preferring ooDSGCs (Huberman et al., 2009; Kay et al., 2011). Due to a genetic linkage between the *Drd4-GFP* transgene and *Flrt2* on chromosome 12, we were only able to obtain three recombinant *Flrt2* mutants carrying the transgene. However, in these cases we observed laminar targeting errors resembling those made by Hb9-DSGCs (Supplementary Fig. 3S5). Thus, *Flrt2* is required to prevent multiple types of ooDSGC dendrites from straying into inappropriate IPL regions.

In addition to these ooDSGC dendritic errors, we also observed a second phenotype in *Flrt2^Ret^* mutants whereby starburst neurons sporadically mistargeted their arbors primarily into the S1 IPL sublayer (Fig. 4A,B,D). In cases that were scored as starburst errors, ooDSGC dendrites were aligned with the mistargeted starburst arbors, consistent with our past findings that ooDSGCs always follow starburst laminar errors (Ray et al., 2018). Therefore, despite being mistargeted, ooDSGC dendrites remained paired with starburst arbors. This is in sharp contrast to the first type of error described above, in which ooDSGCs uncouple from the starburst dendrite scaffold (Fig. 3A-D). Hence, deletion of FLRT2 results in two separate types of laminar errors: one in which pairing between DS circuit partners is disrupted, and a second in which the two cell types make errors together.

**Figure 4:**
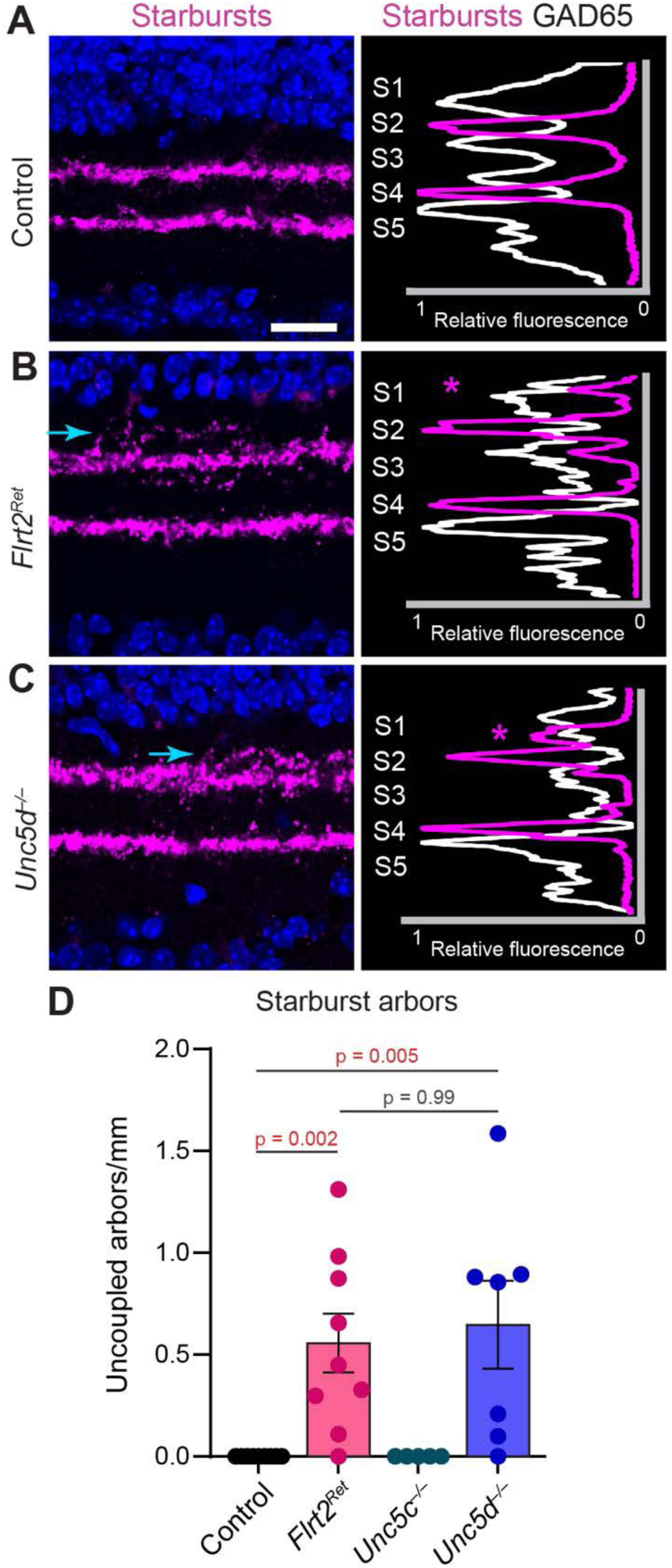
*Flrt2* and *Unc5d* are required for starburst laminar targeting. **A-C**. Representative images (left) and fluorescence profile plots (right) showing starburst amacrine cell IPL projections in retinas of the specified genotype. Magenta, anti-Vacht; white, anti-GAD65. In *Flrt2^Ret^* (B) and *Unc5d^−/−^*(C) animals, starburst dendrites project ectopically to S1 IPL sublayer (arrows; asterisks). Wild-type littermate control starbursts (A) project only to S2 and S4. **D**. Summary of starburst laminar targeting errors, defined as number of VAChT^+^ arbors that were uncoupled from the main starburst scaffold and resided in ectopic sublayers. Mistargeting frequency was similar in *Flrt2^Ret^* and *Unc5d^−/−^* mutants. No targeting errors were observed in *Unc5c^−/−^* mutants (also see Supplementary Fig. 4S2). Statistics, Kruskal-Wallis test (main effect p = 2 x 10^-4^) followed by Dunn’s post-hoc test. P-values (shown on graph) were corrected for multiple comparisons. Red text denotes significant difference. Sample sizes (number of animals): littermate controls n = 9, *Flrt2^Ret^* n = 9, *Unc5c^−/−^* n = 5; *Unc5d^−/−^* n = 7. Error bars, S.E.M.

To ascertain whether starburst laminar targeting errors could result from defects in cross-circuit repulsion, we cultured primary neurons from neonatal retina and measured the overlap between starburst and GAD65^+^ neurites. Overlap was significantly greater in cultures from *Flrt2^Ret^* mutants compared to *Flrt2^WT^* control cultures, suggesting a defect in cell-cell avoidance when FLRT2 is absent (Supplementary Fig. 4S1).

Together, the phenotypes of *Flrt2^Ret^* mutants indicate that FLRT2 influences precise laminar targeting by both starburst cells and ooDSGCs: It prevents starburst dendrites from straying into S1, and it prevents ooDSGC dendrites from departing the starburst scaffold and straying into S1 or S3.

### Cell-type-specific roles for *Unc5c* and *Unc5d* in DS circuit laminar targeting

The second approach we used to disrupt FLRT2-UNC5C signaling was to eliminate UNC5C expression. To this end, we crossed *Hb9-GFP* into an *Unc5c* constitutive mutant background (Ackerman et al., 1997; Kim and Ackerman, 2011) and examined DS circuit IPL stratification at P15-17. Because *Unc5c^−/−^*animals on a pure C57Bl6/J background rarely survived beyond P0, these experiments used a hybrid C57-SJL background (see Methods). We found that mistargeted Hb9-DSGC arbors were rare in *Unc5c^+/+^* control retinas, similar to *Flrt2^WT^* controls. By contrast, there were numerous ooDSGC dendrite targeting errors in *Unc5c^−/−^* mutants (Fig 3A,B,E; Supplementary Fig. 3S6). These ooDSGC dendrite errors strongly resembled those observed in *Flrt2^Ret^* mutants, in that they selectively targeted S1 and S3; moreover, ooDSGC errors occurred at similar frequency in each mutant (Fig. 3A-E; Supplementary Fig. 3S6). These findings support the notion that UNC5C prevents mistargeting of ooDSGC arbors into inappropriate IPL regions via transcellular interactions with FLRT2.

Whereas ooDSGC errors were highly similar in *Flrt2^Ret^* and *Unc5c^−/−^* mice, starburst IPL projections were entirely normal in *Unc5c^−/−^* mutants (Fig. 4D). We therefore considered whether a different FLRT2 binding partner might contribute to starburst laminar restriction. UNC5D is a close homolog of UNC5C that can also bind FLRT2, and which also localizes to non-DS IPL regions in a pattern resembling UNC5C (Visser et al., 2015). Furthermore, similar to *Unc5c*, *Unc5d* is also broadly expressed by non-starburst GABAergic amacrine cells (Supplementary Fig. 2S1). These findings prompted us to examine DS circuit IPL targeting in *Unc5d^−/−^* mutant mice (Yamagishi et al., 2011). Hb9-DSGC dendrite targeting was unaffected by loss of *Unc5d* (n = 3; Supplementary Fig. 4S2). By contrast, a subset of starburst arbors were mistargeted, a phenotype strikingly similar to the one observed in *Flrt2^Ret^* mutants (Fig. 4C,D). Thus, UNC5D appears more important for starburst laminar targeting than UNC5C. Taken together with our results in *Flrt2^Ret^* mutants, these genetic experiments support a model in which precise targeting of ooDSGC dendrites requires UNC5C-FLRT2 signaling, while starburst dendrite targeting mainly involves UNC5D-FLRT2 signals.

### Mistargeted ooDSGC arbors form non-starburst inhibitory synapses

We next investigated the consequences of removing the FLRT2-UNC5C signaling system for DS circuit synaptic specificity. Normally, starburst amacrine cells provide virtually all of the inhibitory synapses onto ooDSGC dendrites (Bleckert et al., 2018; Mani et al., 2022; Sivyer et al., 2019; Yoshida et al., 2001). However, mutant ooDSGCs send dendrites into IPL regions lacking starburst arbors (Fig. 3), raising the possibility that they might receive inhibitory synapses from inappropriate presynaptic partners. To test whether inhibitory synapses are present on mistargeted Hb9-DSGC dendrites, we labeled synapses using antibodies to the presynaptic active zone marker bassoon and the inhibitory post-synaptic marker gephyrin (Fig. 5A,B; Supplementary Fig. 5S1). We then performed unbiased semi-automated identification of paired bassoon-gephyrin synapses on Hb9-GFP^+^ arbors using Object Finder software (Della Santina et al., 2013). We assessed both mutants but were particularly focused on *Unc5c^−/−^* animals because they lack mistargeted starburst arbors (Fig. 4D); thus, in these mutants we can be certain that starburst cells are not the source of synapses onto mistargeted ooDSGC dendrites.

**Figure 5:**
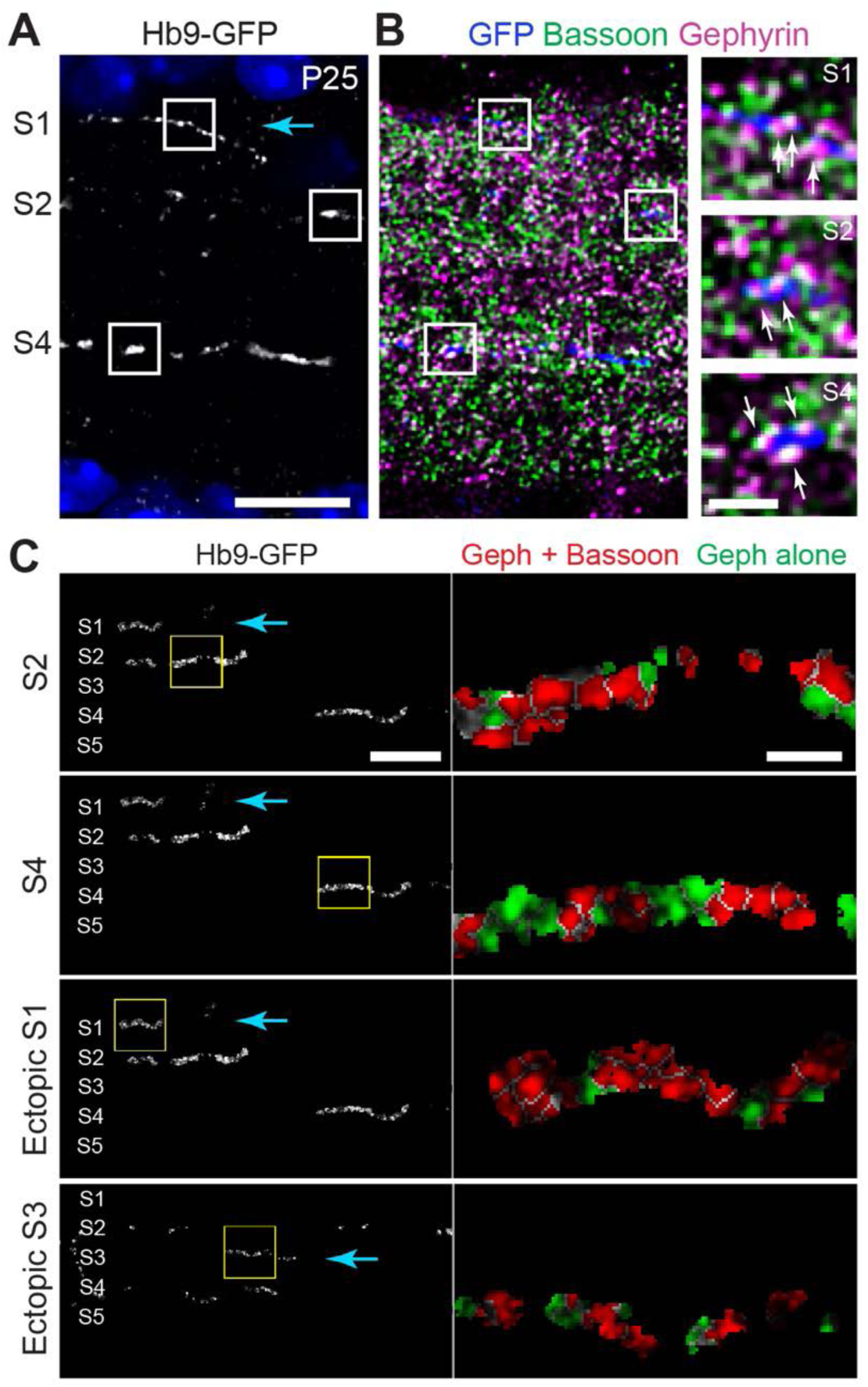
Mistargeted ooDSGC arbors in *Flrt2^Ret^* and *Unc5c* mutants receive inhibitory synapses. **A-B**. Inhibitory synapse labeling in retinal cross-section from P25 *Hb9-GFP*; *Flrt2^Ret^* mutant. Tissue was triple-immunostained for GFP to label ooDSGC dendrites (A); bassoon to label presynaptic active zones (B, green); and gephyrin to label inhibitory postsynaptic sites (B, magenta). All panels show a single optical slice from a representative confocal stack. Boxes denote regions shown at high magnification (B, right). White signals indicate putative synaptic sites where bassoon and gephyrin puncta partially overlap. Putative synapses (arrows) are observed along ooDSGC dendrites regardless of whether they localize to normal DS circuit sublayers (S2, S4) or ectopic sublayers (S1; blue arrow). **C**. Semi-automated unbiased synapse identification in *Hb9-GFP*; *Unc5c^−/−^* retina. ObjectFinder software was used to analyze images similar to B. Left, 3D mask of GFP channel, demarcating ooDSGC dendrites. Right, segmentation of paired bassoon-gephyrin synaptic structures that localize to the GFP mask (red). Green denotes non-synaptic gephyrin signals that were not colocalized with bassoon puncta. Numerous synaptic puncta (red) are evident along ectopic arbors (S1, S3, blue arrows), similar to normally targeted arbors (S2, S4). Images are representative of n = 4 *Unc5c^−/−^* animals and n = 4 *Flrt2^Ret^* animals. Scale bars: 10 µm (A; B, left); 2 µm (B, right) 20 µm (C, left); 5 µm (D, right).

This analysis revealed that mistargeted Hb9-DSGC dendrites in *Unc5c* mutants were studded with putative inhibitory synapses, at a density resembling properly localized arbors within S2 and S4 (Fig. 5C). Indeed, inhibitory synapses were identified on all mistargeted S1 and S3 arbors analyzed (n = 4 *Unc5c*^−/−^, n = 4 *Flrt2^Ret^*; 2 sections per animal). These results suggest that mistargeted Hb9-DSGC dendrites receive numerous inappropriate connections from new inhibitory synaptic partners – probably in both mutants, but certainly in *Unc5c^−/−^*. Co-staining with glutamatergic synapse markers (Vglut1 or bassoon paired with PSD-95) revealed that ectopic arbors also made putative excitatory synapses with bipolar cells, although we could not be certain whether those bipolar synapses arose from inappropriate partners. Together, these results indicate that mistargeted ooDSGC dendrites in *Unc5c* and *Flrt2* mutants receive synapses, and that at least some of the synapses arise from inhibitory non-DS circuit amacrine cells.

### Maintenance of DS circuit laminar targeting mediated by FLRT2-UNC5 repulsion

To understand the mechanism by which FLRT2 and UNC5s control dendritic laminar targeting, we first sought to determine the developmental events that depend on the presence of these molecules. Initial formation of the DS circuit IPL sublayers is a two-step process involving stratification of starburst arbors by P2, and ooDSGC dendrites by P6 (Fig. 6A,B). Both of these events occurred normally in P6 *Flrt2^Ret^* and *Unc5c^−/−^* mutants, indicating that these genes are dispensable for initial circuit assembly (Fig. 6E,F; Supplemental Fig. 6S1).

**Figure 6:**
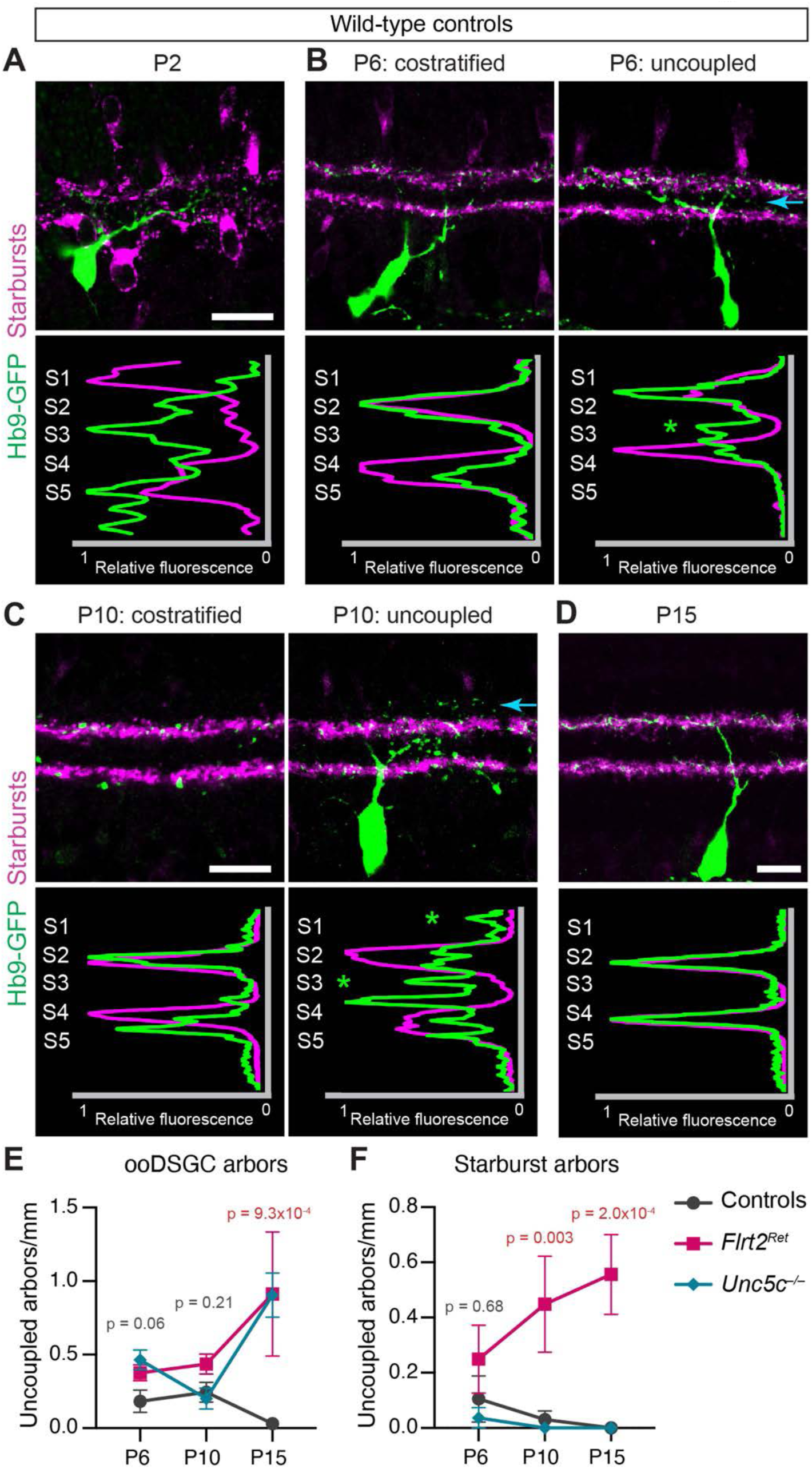
Refinement of DS circuit laminar targeting between P6 and P15. **A-D.** Developmental timecourse showing starburst and Hb9-ooDSGC arborization patterns at the specified developmental ages. All tissue from wild-type animals (i.e. *Flrt2^+/+^; Unc5c^+/+^*); see Figs. 6S1, 6S2 for mutant images. At P2 (A), two starburst IPL strata are evident but Hb9-ooDSGC arbors are not yet stratified. By P6 (B), and continuing at P10 (C), the two cell types are largely co-stratified although uncoupled ooDSGC arbors outside S2/S4 are evident (right panels; arrows). By P15 (D) all ooDSGC dendrites are properly coupled with starburst scaffold. Scale bars, 20 µm (bar in A also applies to B). **E**. Quantification of ooDSGC arbors that are uncoupled from the main starburst scaffold. Data are shown for *Flrt2^Ret^* mutants; *Unc5c^−/−^*mutants; and their wild-type littermate controls. P15 data are replotted from Fig. 3 to facilitate comparisons with younger ages. At both P6 and P10, there was no significant effect of genotype on ooDSGC laminar error rate. Statistics: Kruskal-Wallis test; P-values shown on graph are for main effect of genotype at each age. Sample sizes: P6 control n = 7; P6 *Flrt2^Ret^*n = 8; P6 *Unc5c^−/−^* n = 4; P10 control n = 8; P10 *Flrt2^Ret^*n = 4; P10 *Unc5c^−/−^* n = 6. Error bars, S.E.M. **F**. Quantification of starburst arbors that are uncoupled from the main starburst scaffold. P15 data are replotted from Fig. 4. Statistics: Kruskal-Wallis test; P-values shown on graph are for main effect of genotype at each age. Sample sizes: P6 control n = 7; P6 *Flrt2^Ret^*n = 8; P6 *Unc5c^−/−^* n = 4; P10 control n = 8; P10 *Flrt2^Ret^*n = 5; P10 *Unc5c^−/−^* n = 6.

We next investigated DS circuit laminar development between P6 and P15. This period encompasses the functional maturation of the DS circuit (Tiriac et al., 2022; Wei et al., 2011; Yonehara et al., 2011), and is characterized by horizontal expansion of starburst and ooDSGC arbors within S2 and S4 (Fisher, 1979; Kim et al., 2010; Lefebvre et al., 2012). In wild-type retina, we noticed a previously unreported refinement of ooDSGC lamination that occurred during this time. While the gross stratification pattern of DS circuit arbors was mature by P6, we observed occasional mistargeted ooDSGC dendrites at P6 and P10 that were not paired with the starburst laminar scaffold (Fig. 6B,C). These mistargeted dendrites were eliminated by P15 (Fig. 6D,E). In *Flrt2^Ret^* and *Unc5c^−/−^* mutants, the number of mistargeted ooDSGC dendrites was similar to controls at P6 and P10 (Fig. 6E; Supplementary Fig 6S1). However, instead of eliminating mistargeted dendrites between P10 and P15, both mutants showed a substantial increase in mistargeting during this time (Fig. 6E). A similar phenomenon occurred for starburst arbors between P6 and P10 (Fig. 6F). These findings indicate that, even after targeting the correct IPL sublayer by P6, growing DS circuit dendrites are not confined to the appropriate sublayer but rather continue to produce mistargeted arbors, which are eliminated in a manner that depends on FLRT2-UNC5 signaling.

### UNC5C suppresses elaboration of DS circuit arbors to guide laminar targeting

The results presented thus far support a working model in which FLRT2-UNC5 binding suppresses growth of nascent DS circuit dendrites that have strayed into UNC5^+^ territory (Fig. 7F). If this is true, then misexpressing UNC5C within the DS circuit IPL sublayers should make this region inhospitable to growth of correctly targeted dendrites, thereby favoring growth within ectopic sublayers. To test this prediction, we generated an adeno-associated virus (AAV) construct that expresses HA-tagged UNC5C protein in a Cre-dependent manner (AAV-CMV-flex-UNC5C-HA). This virus was introduced at P1-2 into the retina of *Chat^Cre^* transgenic mice (Fig. 7A), which express Cre recombinase selectively in starburst cells (Ray et al, 2018). This AAV misexpression strategy is denoted Chat-UNC5C. Selective expression of UNC5C protein in starburst cells was validated by anti-HA immunostaining (Supplementary Fig. 7S1). A flex-tdTomato AAV was used as an injection control (Fig. 7A).

**Figure 7:**
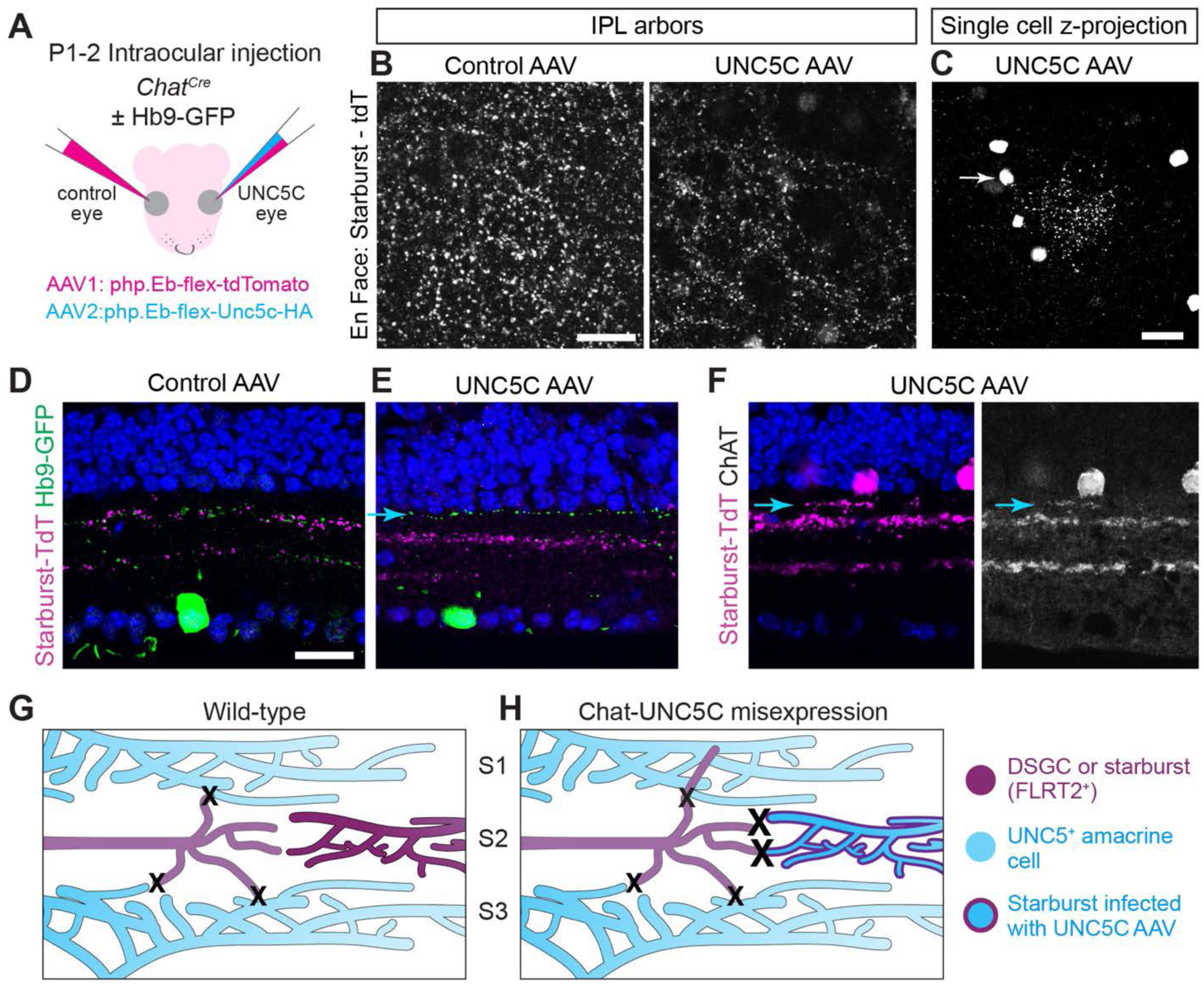
Misexpression of UNC5C in starburst cells induces DS circuit laminar targeting errors. **A**. Schematic showing strategy for starburst-specific AAV expression in *Chat^Cre^* mice. Expression of tdTomato and UNC5C was Cre-dependent due to flex switch. UNC5C eye received both viruses while control eye received only tdTomato. Php.Eb, AAV serotype. **B**. Representative en-face views of retinal wholemounts infected with specified AAVs. Confocal images showing infected starburst arbors were acquired at level of OFF starburst plexus. Images are Z-projections of 3 confocal slices encompassing 1.2 µm near INL/IPL border. In tdTomato-only controls (left, n = 4), starburst dendrites ramify extensively within S2 through transduced retinal region. In UNC5C misexpressing animals, arborization was diminished (n = 5). Also see Supplementary Fig 7S1 which shows density of OFF starburst soma labeling in these regions. **C**. En-face view of a particularly strongly affected region of Chat-UNC5C AAV retina. Image is Z-projection of confcoal stack encompassing INL and IPL, such that both OFF starburst somata and arbors should be visible. Most somata in this field of view make minimal IPL projections. One starburst cell (arrow) with arbors visible in this field of view exhibited highly unusual dendritic morphology. **D-F**. Representative cross-sections showing IPL projections of starburst and ooDSGC dendrites in retinas infected with specified AAVs. Stratification was normal in retinas expressing tdTomato alone (D; n = 4 mice). Chat-UNC5C misexpression caused Hb9-DSGCs (E) and starburst cells (F) to become mistargeted in ectopic IPL layers (starbursts, n = 5 mice; ooDSGCs, n = 4 mice). Green, GFP^+^ Hb9-DSGCs; Magenta, tdTomato^+^ starburst cells; white, anti-Chat starburst staining. **G,H**. Working model of dendrite selection via relative UNC5 signaling. In normal development (G), mistargeted FLRT2^+^ arbors (magenta) are suppressed by contact with UNC5^+^ neurites (blue) within S1 or S3 (small X). By contrast, correctly targeted arbors do not encounter UNC5s, allowing them to survive. H: Upon overexpression of UNC5C within the DS circuit sublayers (e.g. S2), correctly targeted arbors receive a large inhibitory signal (large X), thereby suppressing growth. Mistargeted arbors are able to grow into UNC5C^+^ regions due to relatively lower UNC5 levels (blue shading indicates UNC5 protein levels). Scale bars, 20 µm. Bar in D applies to E,F.

Compared to starburst cells expressing tdTomato alone, starbursts from retinal regions transduced with Chat-UNC5C AAV exhibited altered morphology consistent with impaired elaboration of arbors within sublayer S2 (Fig. 7B; Supplementary Fig. 7S1; n = 5/5 wholemount retinas from 5 Chat-UNC5C mice). Furthermore, we observed starburst laminar targeting errors in Chat-UNC5C animals that were absent from tdTomato-only controls (Fig. 7C,D; n = 4/4 controls without laminar errors; n = 5/5 Chat-UNC5C with errors). These errors closely resembled the starburst phenotype in *Flrt2^Ret^* (Fig. 3) and *Unc5d*^−/−^ mutants (Fig. 4). We also observed mistargeting of Hb9-DSGC dendrites in Chat-UNC5C mice (n = 4/4 controls without errors; n = 4/4 Chat-UNC5C mice with errors), indicating that starburst-derived UNC5C acts in a trans-cellular manner to produce laminar targeting errors (Fig. 7C,E). Altogether, these results indicate that ectopic UNC5C favors elaboration of mistargeted DS circuit arbors at the expense of correctly targeted ones.

As the inappropriate sublayers targeted in Chat-UNC5C mice are also UNC5C^+^, these results demonstrate that UNC5C is not an absolute suppressor of FLRT2^+^ dendrite growth. Instead, DS circuit neurons can elaborate arbors within IPL regions containing endogenous levels of UNC5C when forced to do so by overexpression of UNC5C within S2 and S4. Thus, our findings suggest that DS circuit neurons can select nascent dendrites for further growth based on relative levels of UNC5C signaling (Fig. 7F,G).

### Direct FLRT2-UNC5 protein interactions mediate DS circuit laminar targeting

Finally, we addressed the cellular and molecular mechanisms by which FLRT2 and UNC5s suppress elaboration of nascent dendrites. In addition to UNC5s, FLRT2 can also bind to LPHN-family proteins, which mediate cell adhesion and synaptogenesis upon binding to FLRTs (Jackson et al., 2015; Lu et al., 2015; O’Sullivan et al., 2012; Sando et al., 2019). Transcriptomic studies show that *Lphn1* and *Lphn3* are broadly expressed by most RGCs and amacrine cells, including starburst neurons and ooDSGCs (Kay et al., 2012; Rheaume et al., 2018; Tran et al., 2019; Yan et al., 2020), raising the possibility that FLRT2-LPHN interactions could also contribute to laminar targeting.

To learn the extent to which DS circuit laminar targeting depends on a direct FLRT2-UNC5 protein interaction, we selectively disrupted FLRT2-UNC5 interactions in vivo. Structural studies at atomic resolution led to the identification of a FLRT ectodomain point mutation, FLRT2^H170N^ (Fig. 8A), that blocks UNC5 binding without altering trafficking, surface expression, or interactions with LPHNs (Lu et al., 2015; Seiradake et al., 2014). This so-called FLRT2^UF^ mutation has been used extensively to test functional consequences of FLRT2-UNC5 signaling (Fleitas et al., 2021; Jackson et al., 2015; Jackson et al., 2016; Lu et al., 2015; Seiradake et al., 2014). Thus, FLRT2^UF^ provides a useful genetic tool to test whether FLRT2-UNC5 binding is required for DS circuit laminar targeting. Using CRISPR editing, we introduced the *Flrt2^UF^* point mutation into the mouse germline. *Flrt2^UF/UF^* animals (abbreviated *Flrt2^UF^*) on a pure C57Bl6 background rarely survived to P0. However, similar to *Unc5c^−/−^*mice, *Flrt2^UF^* mutants did survive and appeared grossly normal when bred on a mixed C57Bl6-SJL background. Therefore, we used this mixed background to investigate postnatal retinal phenotypes in *Flrt2^UF^* mice.

**Figure 8:**
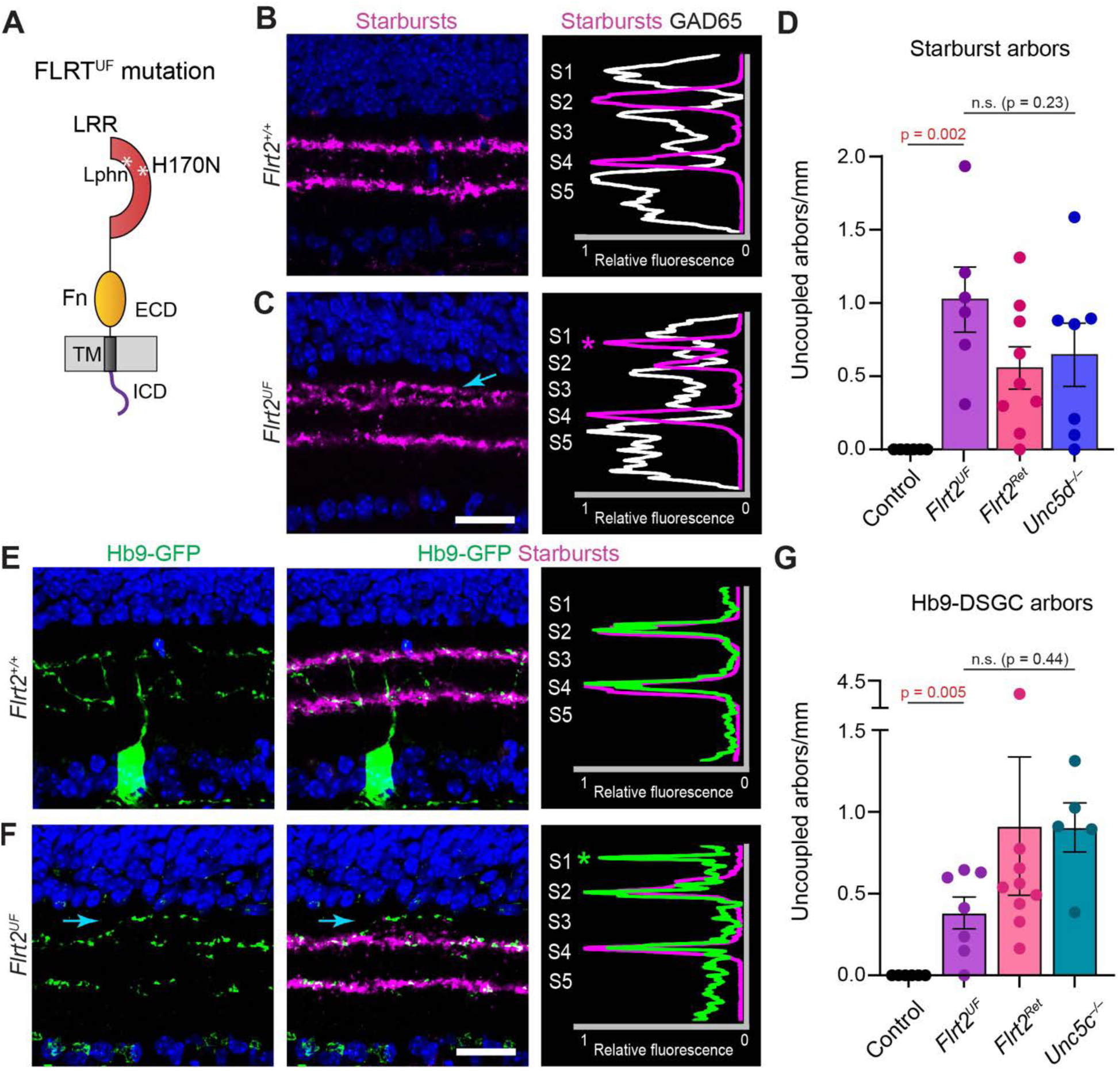
Abolishing FLRT2-UNC5 binding disrupts DS circuit laminar targeting. **A**. Schematic of FLRT2^UF^ mutant protein. H170N mutation introduces a N-glycosylation site within leucine-rich repeat (LRR) domain that interferes with the UNC5 binding surface. LPHN binding site is unaffected. TM, transmembrane domain; ECD, extracellular domain; ICD, intracellular domain. **B,C.** Effects of *Flrt2^UF^* mutation on starburst dendrite targeting. Left panels, VAChT immunostaining; right panels, profile plots. In littermate control *Flrt2^+/+^* retinas (B), starbursts occupy S2 and S4, avoiding regions with GAD65^+^ arbors. In homozygous *Flrt2^UF^* mutants (C), starbursts make laminar targeting errors into S1 (C, arrow), reminiscent of the *Flrt2^Ret^* and *Unc5d^−/−^*phenotype. These arbors invade GAD65 territory (right panel, asterisk). **D**. Number of mistargeted starburst arbors in *Flrt2^UF^* mutants and *Flrt2^+/+^* littermate controls. Statistics, two-tailed Mann-Whitney test; sample sizes n = 6 littermate controls; n = 6 *Flrt2^UF^*mutants. For comparison, *Flrt2^Ret^* and *Unc5d^−/−^* mutant data are replotted from Fig. 4. There was no significant difference in error frequency between *Flrt2^UF^* and the other two mutants (one-way ANOVA, F(2, 19) = 1.60; n.s., not significant). Statistics: Kruskal-Wallace test (main effect p = 0.003) with post-hoc Dunn’s test corrected for multiple comparisons. P-values shown on graph. Sample sizes: n = 6 littermate controls; n = 6 *Flrt2^UF^* mutants; n = 7 *Unc5d^−/−^* (see Fig. 4); n = 9 *Flrt2^Ret^* (see Fig. 4). **E,F.** ooDSGC dendrite targeting in *Flrt2^UF^* mutants (F) and littermate controls (E). Mutant ooDSGCs make laminar targeting errors into S1 (arrow), similar to *Flrt2^Ret^* and *Unc5c* mutants. Control dendrites stratify properly in S2 and S4. Scale bar, 20 µm. **G**. Number of mistargeted Hb9-ooDSGC arbors in *Flrt2^UF^* mutants and *Flrt2^+/+^* littermate controls. Statistics, two-tailed Mann-Whitney test; sample sizes n = 6 littermate controls; n = 7 *Flrt2^UF^* mutants. For comparison, *Flrt2^Ret^* and *Unc5d^−/−^* mutant data are replotted from Fig. 3. There is a trend towards fewer errors in *Flrt2^UF^* compared to the other two mutants but the difference was not significant (one-way ANOVA, F(2, 18) = 0.85). Scale bars, 20 µm. Error bars (D,G), S.E.M.

This analysis revealed mistargeted starburst arbors within the S1 sublayer in *Flrt2^UF^* animals, similar to *Flrt2^Ret^*and *Unc5d*^−/−^ mutants (Fig. 8B-D). Hb9-DSGC dendrites also misprojected into S1 and S3, similar to *Flrt2^Ret^* and *Unc5c^−/−^*(Fig. 8E-G). Comparing *Flrt2^UF^* to *Flrt2^Ret^* mutants, which completely lack retinal *Flrt2* function, the number of starburst errors was comparable (Fig. 8D). Thus, loss of UNC5 binding appears to fully account for the *Flrt2* null mutant phenotype in starburst cells. We also failed to detect a significant difference in ooDSGC mistargeting between *Flrt2^UF^* and *Flrt2^Ret^*; however, we did note a trend towards fewer errors in *Flrt2^UF^*, raising the possibility that other FLRT2 ligands may cooperate with UNC5s to mediate ooDSGC laminar targeting (Fig. 8G). Nevertheless, these findings support the conclusion that UNC5s are the major FLRT2 ligand mediating suppression of mistargeted DS circuit arbors.

### UNC5s influence dendrite targeting by interfering with FLRT-LPHN adhesion

How does FLRT2-UNC5 binding suppress dendrite elaboration? We considered two main possibilities. First, binding might initiate a negative signal, transduced either by FLRT2 or a co-receptor (Leyva-Díaz et al., 2014), that inhibits arbor growth. Alternatively, UNC5 binding might suppress arborization by blocking an affirmative FLRT2-mediated signal, thereby preventing FLRT2 from stabilizing mistargeted arbors. This latter scenario is plausible because FLRT-LPHN adhesion is a well-characterized affirmative signaling system; and the widespread expression of *Lphn1* and *Lphn3* by inner retinal neurons implies that LPHNs should be available throughout the IPL for FLRT2 binding. Furthermore, FLRT2 can bind simultaneously to LPHNs and UNC5s, raising the possibility that UNC5s could influence FLRT-LPHN function (Jackson et al., 2016). We therefore considered the possibility that UNC5s act by occluding FLRT2-LPHN adhesion.

We began by testing the first model – that FLRT2 is part of a receptor complex that transduces repulsive signals upon binding to UNC5s. If this is true, FLRT2 should be required cell autonomously for repulsion. To test this possibility we generated *Flrt2^RGC^* mutant mice in which *Flrt2* was deleted selectively from RGCs. We found that ooDSGC laminar targeting was normal in *Flrt2^RGC^* mutants, indicating that FLRT2 is not required cell autonomously and thus is unlikely to function as a repulsive receptor (Fig. 9A,B).

**Figure 9:**
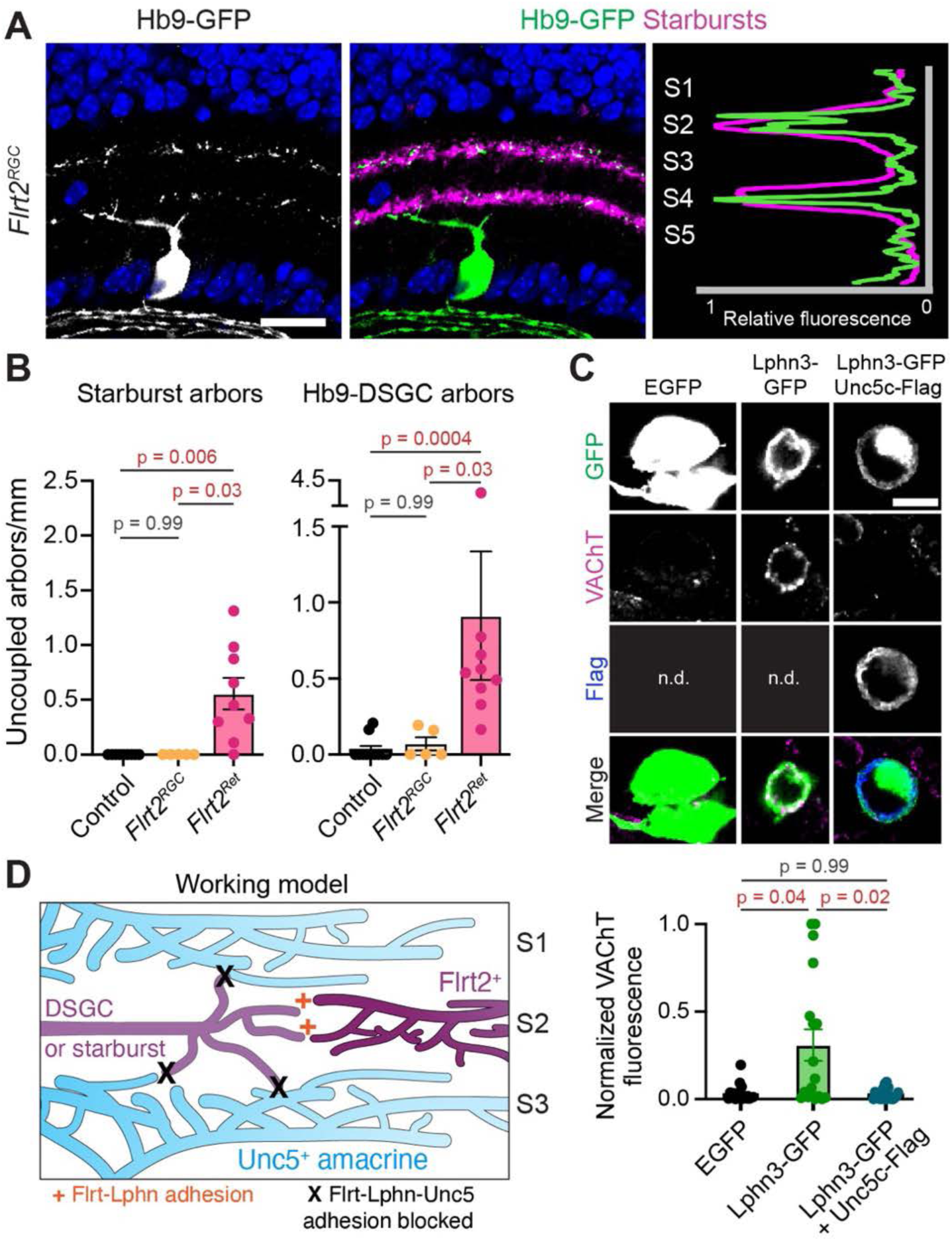
UNC5s function by occluding FLRT-LPHN adhesion. **A,B**. Deletion of *Flrt2* from retinal ganglion cells (*Flrt2^RGC^*) does not cause laminar targeting defects. A, representative image and profile plot from mutant retina. Both ooDSGCs (anti-GFP) and starbursts (anti-VAChT) stratify normally. B, quantification of errors. Sample sizes: controls, n = 9; *Flrt2^RGC^*n = 5. For comparison, *Flrt2^Ret^* data are replotted from Fig. 4D and 3E. Statistics, Kruskal-Wallace test with post-hoc Dunn’s test. Scale bar = 20 um. **C.** Adhesion assay using cultured retinal neurons. Starbursts arbors (marked by anti-VAChT) encircle HEK cells transfected with LPHN3-GFP, but not GFP alone. Co-transfection of UNC5C-Flag blocks the LPHN3 adhesion effect. Top, representative images. Note that cells from left, center columns were not stained for anti-Flag (n.d.) because they were not transfected with Flag construct. Bottom, quantification of VAChT fluorescence surrounding HEK cells transfected as indicated. Sample sizes: n = 18 EGFP; n = 18 Lphn3; n = 20 Lphn3+Unc5c. N = two separate cultures. Statistics, Kruskal-Wallace test followed by Dunn’s post-hoc test. **D**. Working model for role of FLRT2-UNC5C in laminar targeting. FLRT2^+^ dendrites within DS circuit sublayers send out exploratory branches as they grow. Some branches remain properly stratified; these are stabilized by FLRT-LPHN adhesion (+). Other branches are less precisely targeted; they contact UNC5^+^ amacrine arbors in adjacent layers which blocks FLRT-LPHN adhesion, leading to branch elimination (X). Scale bars, 20 µm (A); 10 µm (C). Error bars, S.E.M.

We therefore proceeded to test the second model – that FLRT2-UNC5 binding influences dendrite targeting by occluding FLRT2-LPHN adhesion. This model can only be true if LPHNs are affirmative cues that stabilize nascent DS circuit arbors. To determine whether this is the case, we performed a cell adhesion assay in which primary retinal neurons were cultured with HEK 293 cells expressing LPHN3. Antibodies to the starburst marker VAChT revealed that starburst arbors associated extensively with LPHN3-GFP^+^ HEK cells but not with HEK cells expressing GFP alone (Fig. 9C). Therefore, FLRT2^+^ starburst cells are subject to LPHN3-mediated adhesion, which functions to stabilize starburst arbors that have contacted LPHN3^+^ cells. To test whether UNC5s can interfere with this adhesion, we co-transfected HEK cells with constructs encoding UNC5C and LPHN3. Remarkably, co-expression of UNC5C entirely blocked starburst arbors from adhering to LPHN3 HEK cells (Fig. 9C). This finding indicates that FLRT2-UNC5 binding inhibits FLRT2-LPHN adhesion, and that this anti-adhesive effect is sufficient to influence starburst dendrite targeting.

Taken together with our in vivo results, these observations support a model whereby FLRT2-UNC5 binding supports laminar specificity by diminishing adhesion of mistargeted dendrites, thereby promoting their removal (Fig. 9D).

## DISCUSSION

Rejection of inappropriate synaptic partners is a key step in the establishment of synaptic specificity. It has previously been argued that affirmative cues are sufficient to specify these partner choices: Inappropriate partners may be rejected simply because they lack the correct adhesive code (Lefebvre et al., 2015; Sanes and Zipursky, 2010). Here, we demonstrate that negative cues also contribute in critical ways to laminar and synaptic specificity in the retinal circuit that detects image motion. We show that a direct interaction occurs between FLRT2, which is expressed by DS circuit neurons, and UNC5C/ UNC5D, which are expressed by non-DS GABAergic amacrine cell dendrites stratified outside the DS circuit. This FLRT2-UNC5 interaction is necessary to restrict formation of mistargeted DS circuit arbors. When FLRT2-UNC5 transcellular signals are disrupted, ooDSGCs generate ectopic dendrites that are uncoupled from their starburst partners but which nevertheless acquire numerous inhibitory synapses. We show that the FLRT2-UNC5 system functions during the second postnatal week, after initial establishment of the DS circuit but during a period of growth and refinement of DSGC dendrites. Finally, we provide evidence that the molecular mechanism underlying DS circuit arbor refinement involves the FLRT2-LPHN-UNC5 ternary complex. Together, these findings support a model in which the inhibitory FLRT2-UNC5 system operates in concert with adhesive/attractive molecules, which also play critical roles in DS circuit specificity by providing positive cues that affirm correctly targeted projections (Hamilton et al., 2021). Indeed, our data suggest that FLRT2-LPHN binding is part of the adhesive system that affirms DS circuit contacts, and that UNC5s exert their inhibitory effects by interfering with this adhesion. Thus, our study suggests a mechanism by which inhibitory cues such as UNC5s cooperate with affirmative cues to mediate the rejection of inappropriate partners (Fig 9D).

### FLRT2-UNC5 signaling maintains laminar specificity during dendrite arbor expansion

Our studies of DS circuit IPL development have identified a previously unappreciated developmental phase during which laminar targeting and synaptic specificity are labile. Starburst and ooDSGC IPL associations appear largely adult-like by ∼P6, so prior studies have presumed that laminar targeting is complete by this age (Kim et al., 2010; Peng et al., 2017). However, we show here that FLRT2-UNC5 signaling is required to maintain DS circuit laminar specificity between P6 and P15. In the absence of FLRT2 or its UNC5 ligands, DS circuit neurons stratify properly by P6 but subsequently produce mistargeted arbors. Therefore, the FLRT2-UNC5 system is required to prevent correctly targeted arbors from departing the DS circuit scaffold and straying into neighboring IPL strata. While mistargeted arbors are a normal feature of DS circuit development during the second postnatal week, such arbors are normally eliminated (Fig. 6). By contrast, in mutants these mistargeted arbors remain and elaborate further. Together these observations suggest that developmental events occurring during this period tend to produce mistargeted arbors, which are eliminated by P15 in a FLRT2-UNC5 dependent manner.

What developmental events might cause DS circuit dendrites to wander from their laminar scaffold? At P6, starburst and ooDSGC dendritic arbors are far smaller than they will ultimately be at maturity. During the second postnatal week, both cell types grow substantially in the tangential plane of the retina, reaching their adult arbor sizes by ∼P14 (El-Quessny et al., 2020; Kim et al., 2010; Lefebvre et al., 2012). While it has generally been assumed that such growth can remain tightly associated with the sublayer that was chosen at P6, this may not in fact be possible given that dendrites typically grow by an iterative process in which the stable portion of an arbor sprouts numerous transient processes (Wong and Ghosh, 2002). Only a small subset of these sprouts will survive to become stably incorporated into the mature arbor (Cline and Haas, 2008; Lohmann et al., 2002; Niell et al., 2004; Takeo et al., 2021). Live imaging of starburst cells during the second postnatal week indicates that they too grow their dendrites in this manner (Ing-Esteves and Lefebvre, 2021). Given this growth mechanism, it is plausible that ooDSGCs and starburst dendrites cannot expand in the tangential plane of the retina without inadvertently sprouting into adjacent sublayers. In this case, the FLRT2-UNC5-LPHN system could serve the crucial role of distinguishing correctly targeted dendritic sprouts from mistargeted ones.

Our data support a model (Fig. 9D) whereby the distinction between correct and mistargeted dendritic branches is mediated by relative levels of FLRT2-LPHN adhesion. Such adhesion could in principle occur anywhere throughout the IPL, given that most RGCs and amacrine cells express *Lphn1* and/or *Lphn3* (Kay et al., 2012; Rheaume et al., 2018; Tran et al., 2019; Yan et al., 2020). However, laminar restriction of inhibitory UNC5 proteins creates an adhesion differential that guides dendrite targeting. During the sprouting phase of dendrite growth, ooDSGC and starburst arbors that contact the correct IPL sublayer should experience stronger FLRT2-LPHN adhesion than those contacting adjacent layers, where UNC5s are present and can interfere with such adhesion (Fig. 9D). Branches that experience more FLRT2-mediated adhesion would then be more likely to survive and become part of the stable dendrite tree.

This model is appealing because it explains the DS circuit laminar targeting phenotypes we observed in our gain- and loss-of-function experiments. Under normal circumstances, the adhesion differential between correct and mistargeted arbors may be so great that the chance of mistargeted arbors surviving during the pruning phase is close to zero. However, when UNC5s are absent or when FLRT2^+^ arbors cannot bind UNC5s due to the *Flrt2^UF^* mutation, mistargeted arbors are expected to experience greater FLRT2-LPHN adhesion allowing at least some of them to survive. Conversely, misexpression of UNC5C inside DS circuit sublayers is expected to weaken within-layer FLRT2-LPHN adhesion, thereby promoting elimination of nascent arbors that should have survived. Indeed, UNC5C misexpression caused decreased starburst arbor elaboration and occasional disruptions to the characteristic “starburst” dendritic pattern for which these cells are named. These phenotypes are precisely what would be expected if ectopic UNC5C can drive elimination of correctly targeted dendritic sprouts. Additionally, by weakening the adhesion differential between correct and mistargeted sprouts, UNC5C misexpression and FLRT2 deletion are both expected to increase the chances that mistargeted branches survive to maturity – a phenotype we indeed observed in both *Flrt2^Ret^* mutants and in UNC5C misexpressing animals (Fig. 7).

How might an adhesion differential promote survival of particular branches? FLRT-LPHN interactions are known to mediate synapse formation (O’Sullivan et al., 2012; Sando et al., 2019); this could be the mechanism by which certain branches are selected for stabilization (Cline and Haas, 2008; Niell et al., 2004). However, adhesive interactions can also preserve nascent dendrite branches even in the absence of synaptogenesis (Lohmann and Bonhoeffer, 2008), so it is premature to conclude that synaptogenesis is involved in this process. Further work will be needed to understand the cellular mechanism by which FLRT2 acts to preserve nascent dendrite branches, and how UNC5-FLRT2 binding inhibits this process.

### Implications for wiring of the retinal direction-selective circuit

The retinal DS circuit has been an important model system for understanding mechanisms underlying laminar and synaptic specificity (Hamilton et al., 2021; Prigge and Kay, 2018). Previous work has focused on the P0-P6 period when the DS circuit IPL layers are established. This initial phase of circuit assembly, in which starburst dendrites coalesce to form new IPL layers and then recruit ooDSGC dendrites to join them, requires adhesion molecules like contactins and cadherins (Duan et al., 2014; Duan et al., 2018; Peng et al., 2017). Here we make the unexpected finding that there is a second, later phase of DS circuit laminar targeting, from P6-P15, which uses entirely distinct mechanisms – i.e. FLRT2-UNC5 binding – to ensure that synaptic specificity is maintained. The specific amacrine cell type(s) that synapse onto mistargeted ooDSGC dendrites in *Flrt2* and *Unc5c* mutants remains unclear. However, it is highly unlikely that these synapses are derived from a cell type that normally connects to ooDSGCs. Aside from starburst cells, ooDSGCs can also receive synapses from several other amacrine types, but these synapses are either glutamatergic (Mani et al., 2022) or are targeted to the cell body rather than the dendritic arbor (Bleckert et al., 2018). Thus, the presence of non-starburst inhibitory inputs onto ooDSGC dendrites represents a striking loss of specificity.

It is important to note that only a subset of starburst cells and ooDSGCs made persistent laminar errors in the absence of FLRT2-UNC5 signaling. At first glance this observation appears to suggest that the P0-P6 adhesion-based system has bigger effects on DS circuit anatomy: Widespread ooDSGC errors were observed upon elimination of the cadherins that mediate pairing of ooDSGCs to starburst cells (Duan et al., 2018). However, to completely disrupt the cadherin system, it was necessary to simultaneously knock out three cadherin genes – *Cdh6, Cdh9,* and *Cdh10*. When only a single one of these cadherins is deleted, ooDSGC phenotypes are quite mild – notably milder than the phenotype of *Flrt2* and *Unc5c* single mutants (Duan et al., 2018). It is therefore possible that a more thorough disruption of the FLRT-UNC5-LPHN system would produce more profound phenotypes, particularly since *Flrt3* is also expressed by a subset of DSGCs (Goetz et al., 2022; Ruff et al., 2021; Shekhar et al., 2022). Further work will be needed to test this possibility and thereby to assess the relative importance of the early vs. late laminar targeting systems.

Genetic data suggest a surprising cell type specificity in the function of the two UNC5 proteins: UNC5C appears most important for ooDSGC dendrite targeting, whereas starburst targeting relies more on UNC5D. However, because of their similar expression patterns, it is possible that single mutants do not reveal the full spectrum of UNC5 gene functions. For example, continued presence of UNC5C could occlude ooDSGC phenotypes in *Unc5d* mutants – and likewise for starburst cells in *Unc5c* mutants. Unfortunately we could not test this hypothesis directly because *Unc5c^−/−^; Unc5d^−/−^* double mutants did not survive to birth.

There was also a surprising specificity to the IPL targeting errors induced by manipulation of *Flrt2* and *Unc5* genes. Uncoupled ooDSGC arbors projected selectively to S1 and S3, while starburst errors selectively targeted S1. Furthermore, only the OFF starburst population exhibited errors; ON starburst cells projected normally in both gain- and loss-of-function experiments. The reasons why ON starbursts were spared remain unclear; one possibility is that these cells are subject to extra adhesion, mediated by the ON starburst-specific molecule contactin-5 (Peng et al., 2017), and are therefore less reliant on repulsive mechanisms for their laminar targeting. In considering why mistargeted arbors might stratify within specific sublaminae, we noted that the IPL strata which generally lacked ectopic arbors – between S4 and the GCL – is also the region containing the specialized circuitry of the rod pathway. Perhaps this IPL territory, which is dominated by arbors from rod bipolar cells and AII amacrine cells, is not permissive for survival of mistargeted ooDSGC or starburst dendritic branches. Another possible explanation for the bias towards S1 and S3 is that mistargeted starburst and/or ooDSGC dendrites might encounter affirmative molecular cues within these strata that stabilize errant arbors. Given this possibility, it is notable that we previously found two other members of the FLRT protein family, FLRT1 and FLRT3, to be strongly expressed within S1 and S3 respectively (Visser et al., 2015). Each of these FLRT family molecules can bind to LPHNs to mediate adhesion (O’Sullivan et al., 2012); however, it remains to be determined whether such adhesion has a role in guiding mistargeted arbors. Further experiments probing for additional DS circuit stabilizing cues will be needed to fully understand how mistargeted arbors choose their alternate targets.

### New functions and mechanisms for FLRT-UNC5 signaling

FLRT and UNC5 proteins are emerging as important contributors to many facets of brain development and disease (Ando et al., 2022; Del Toro et al., 2017; Khani et al., 2022; Klein and Pasterkamp, 2021; Murcia-Belmonte et al., 2019; Yamada et al., 2019). Thus, there has been substantial interest in understanding both the molecular mechanisms by which these proteins function as well as the specific neurodevelopmental events they control. Previous studies of FLRT-UNC5 receptor-ligand interactions have emphasized effects on UNC5-expressing cells, mediated by UNC5s in their capacity as repulsive guidance receptors (Fleitas et al., 2021; Yamagishi et al., 2011). By contrast, our results reveal that signaling in the reverse direction is also possible: We show that FLRT2-expressing dendrites can be sculpted by UNC5 binding. This signaling configuration had been suggested previously (Visser et al., 2015) but had not been shown to operate in vivo. The mechanism by which UNC5-to-FLRT signaling sculpts dendrites is somewhat unconventional: Rather than using FLRT2 as a repulsive receptor or co-receptor (Leyva-Díaz et al., 2014), UNC5 binding appears to guide dendrites by modulating FLRT adhesion. Our data raise the intriguing possibility that FLRT2-UNC5 binding may mediate bidirectional repulsion – a mechanism that would have obvious utility in establishing discrete IPL sublayers. In this study we did not thoroughly investigate whether stratification of UNC5^+^ amacrine cells is affected by loss of FLRT2. Although no obvious phenotypes were evident using GAD65 immunostaining, it is possible that analysis of specific UNC5C- or UNC5D-expressing populations would reveal targeting errors. Such studies await improved reagents for selective labeling of UNC5C^+^ or UNC5D^+^ amacrine populations.

Many previous studies have shown that FLRTs bind to LPHNs in vivo. This interaction promotes transcellular adhesion and synaptogenesis, as noted above, as well as cell migration (del Toro et al., 2020; O’Sullivan et al., 2012; Sando et al., 2019; Schroeder et al., 2018). While it was known that UNC5s can form a ternary complex with LPHNs and FLRTs, the function of the complex was not known. Here we provide evidence that presence of UNC5s within this complex inhibits FLRT-LPHN adhesion to guide dendrite growth in vivo. This inhibitory function of UNC5 proteins likely involves forming a tripartite complex with FLRTs and LPHNs because 1) inhibition requires FLRT2 binding; 2) LPHNs are broadly expressed and likely to be available for binding throughout the IPL. However, it is also possible that UNC5 binding sequesters FLRT2 to prevent it from engaging in adhesive LPHN interactions. Either way, it is clear from our work that UNC5 binding is a key modulator of FLRT2 adhesive functions, with crucial effects upon dendrite elaboration and synaptic specificity. Therefore, due to the widespread expression of these molecules, the mechanism we have described here has the potential to be broadly relevant to dendrite development and synaptic partner choices throughout the nervous system.

## METHODS

### Mice

Mice of both sexes were used for experiments. CD1 mice were obtained from Charles River laboratory. *Flrt2^flox^* mice (*Flrt2^tm1c(EUCOMM)Wtsi^/RobH*) were obtained from the EMMA repository (EM: 08315). Mice carrying the *Unc5c^rcmTg(Ucp)1.23Kz^* null allele (Ackerman et al., 1997) were a gift of Dr. Susan Ackerman (UCSD). *Unc5d^tm1Kln^*null mice (Yamagishi et al., 2011) were a gift of Dr. Victor Tarabykin (Charité, Berlin) and were kindly provided by Dr. David Feldheim (UCSC). Drd4-GFP (*Tg(Drd4-EGFP)W18Gsat*)) mice were a gift of Dr. Joshua Sanes. *Flrt2^UF^* mice were generated in this study as described in detail below. The following strains were obtained from Jackson Labs – see Key Resources table for catalog numbers: Hb9-GFP (*B6.Cg-Tg(Hlxb9-GFP)1Tmj/J*); Six3-Cre (*Tg(Six3-cre)69Frty*)); *Chat^Cre^* (*Chat^tm2^*^(*cre)Lowl*^/*J*); Vglut2*^Cre^ (Slc17a6^tm2(cre)Lowl^*/J).

Mutant and transgenic strains were maintained on the C57Bl6/J background, and in most cases experimental animals were also on this background. For *Unc5c* null mice, experimental animals were generated on a mixed C57-SJL background as follows: First, to generate breeders for experimental matings, heterozygous mutants (*Unc5c^+/–^*) on the C57Bl6 background were outcrossed to SJL/J. *Unc5c^+/–^* progeny from this cross were then bred to *Unc5c^+/–^* or Hb9-GFP; *Unc5c^+/–^*animals on the pure C57Bl6 background, thereby generating mutants and wild-type littermate controls for experiments. A similar breeding strategy was used for the *Flrt2^UF^* strain.

To generate *Flrt2^Ret^* mutants, *Six3-Cre; Flrt2^flox/+^* mice were bred to *Flrt2^flox/+^* or *Flrt2^flox/flox^*animals with or without the Hb9-GFP transgenes. These crosses yielded *Flrt2^Ret^*mutant animals of genotype *Six3-Cre; Flrt2^flox/flox^*, as well as two types of wild-type controls 1) *Six3-Cre; Flrt2^+/+^* and 2) *flox/+* or *flox/flox* animals without the Cre transgene. These were phenotypically indistinguishable and so were pooled as *Flrt2^WT^* animals. A similar breeding strategy was used with Vglut2^Cre^ mice to generate *Flrt2^RGC^*mutants.

For all strains, P1-P6 neonates were euthanized by ice anesthesia followed by decapitation. Mice aged P7 and older were euthanized via isoflurane anesthesia followed by decapitation.

### Generation of *Flrt2^UF^* mice

To engineer the *Flrt2^UF^* mutation (H170N; Seiradake et al., 2014) into the mouse genome, we used a CRISPR-Cas9 homology-directed repair strategy. We generated a 200 bp repair oligonucleotide homologous to the region surrounding the codon encoding amino acid H170. In this oligonucleotide the CAC H170 codon was mutated to AAC (N). To induce homology-directed repair at the appropriate genomic site, a single guide RNA (sgRNA) was produced in which the PAM sequence overlaps the H170 codon (sequence 5’ GCCCAACAGGCACGCTACTCagg; PAM site is denoted by lowercase letters). The PAM sequence is destroyed by the codon-altering mutation, thereby rendering correctly edited alleles immune to further Cas9 cutting activity. See Key Resources table for sequences.

The repair DNA was electroporated into one-cell stage embryos together with pre-assembled Cas9-sgRNA ribonucleoproteins, as described (Modzelewski et al., 2018). Briefly, to generate sgRNA for electroporation, a DNA template was generated by PCR and then used for RNA synthesis (HiScribe T7 RNA polymerase, NEB #E2040S). DNA template was removed from the reaction with Turbo DNAse (Invitrogen #AM2238). The sgRNA was purified by phenol/chloroform extraction, ethanol precipitated, and resuspended in nuclease-free H_2_O. The Cas9 ribonucleoprotein was assembled by incubating Cas9 protein, sgRNA, and repair oligo together for 10 min at 37°C in Tris (2-carboxyethyl) phosphine hydrochloride (TCEP) buffer (Modzelewski et al., 2018). The ribonucleoprotein mix was electroporated using a NEPAGENE Super Electroporator (NEPA21 Type II) using a glass slide chamber with platinum electrodes (Bulldog Bio part CUY501P1-1.5) with the following settings: Poring Pulse: V 40, Length(ms) 3.5, Interval (ms)50, No 4, D. Rate % 10, Polarity +; Transfer Pulse: V 7, length (ms) 50, Interval (ms) 50, No. 5, D. Rate% 40, Polarity +/-. Following electroporation the embryos were transferred to pseudopregnant females.

To screen founders for the correctly modified allele, founder genomic DNA was prepared from tail and/or toe tissue and subject to PCR using primers targeting the ends of the repair oligonucleotide homology region (Sense: ‘5’ accgcggtggcggcc GCT CCT CAA GCT GGA AGA ACT CCA 3’; Antisense: 5’ tagaggatccactag TGT CTG ATA TGA CGG CAA TTC GGT 3’). PCR products were subject to Sanger sequencing using Sense primer. For animals in which sequencing revealed InDels across the target region, the PCR fragments containing InDels were cloned into pL253 at the NotI and SpeI sites via InFusion cloning (Takara #ST0344). Bacterial recombinants were screened via PCR using primers 253.S (caaggcgattaagttgggtaac) and 253.AS (gcgtgttcgaattcgccaatgac). Plasmid DNA from positive clones were sequenced using Sense primer for verification of the targeted mutation. Positive founders were outcrossed to C57Bl6/J.

To genotype the UF dinucleotide point mutation by PCR, we used a 4 primer strategy (Peng et al., 2018) in which “outer” primers in constant regions combine with “inner” primers overlapping the point mutation site to distinguish the wild-type and mutant alleles. Primer sequences are given in the Key Resources table. Band sizes: Outer, 599 bp. Inner mutant, 426 bp. Inner wild-type, 231 bp.

### Cell culture

Primary cultures of retinal neurons were prepared using P0/P1 mice obtained from CD1 timed pregnant litters (Charles River). The procedure was modified from the protocol of Jiang et al. (2019). Retinas were isolated from the eyecup in dissecting solution (Jiang et al., 2019), followed by incubation in 0.5% Trypsin for 15-20 min at 37°C with gentle intermittent mixing. Next, trypsin was exchanged two times with dissecting solution and reaction was stopped using stopping solution (HBSS + 5% serum). Trypsinized retina was triturated 4 times either in stopping solution or MEM1X solution and centrifuged for 5 mins at 800 x g at 4°C. Cells were resuspended in 1.0 ml of plating medium (Jiang et al., 2019) and filtered through 30 µm sterile filter. Isolated cells were plated at density of ≥ 200,000 cells/well on previously coated Poly L-lysine coverslips. Cells were allowed to settle for 30 mins in cell culture incubator, at which time 2 ml of plating medium was added into each well. Cells were incubated for 1 day at 37°C. On the next day, media was exchanged for neuronal growth medium (Jiang et al., 2019) containing B27 supplement. On day 4, AraC (cytosine arabinoside) was included in the neuronal growth medium. Cultures were fixed on day 10-12.

For GAD65-VAChT, overlap analysis, confocal stacks were acquired from GAD65 + VAChT immunostained cultures and single z slices chosen for analysis. Overlap analysis (Manders et al., 1992) was performed with coloc2 ImageJ plugin. The overlap ratio was calculated as the VAChT pixel intensities within pixels containing non-zero GAD65 values, divided by total VAChT pixel intensity across the image. This yields a measure of overlap between the two signals that is independent of overall image brightness (Manders et al., 1992).

For the cell adhesion assay, primary cultures were prepared as described above, and transfected HEK 293 cells were added to the culture on day 8. The day prior to co-culture, HEK cells were transfected with the following plasmids: CMV-EGFP; CMV-Lphn3-GFP encoding full-length LPHN3 (O’Sullivan et al., 2012); and/or CMV-Unc5c-Flag encoding full-length UNC5C (Visser et al., 2015). Transfected cells were co-cultured with primary cells for 2 days, then fixed in 2% paraformaldehyde/1x PBS and subjected to immunohistochemistry.

### RNAScope In Situ Hybridization

Eyes from euthanized P6 wildtype mice were fixed in 4% paraformaldehyde in PBS for 24 hours at RT and then washed 3X with PBS. Isolated retinas were cryoprotected in 30% sucrose and 0.02% sodium azide in PBS and frozen in Tissue Freezing Medium (General Data; Cincinnati, OH). Retinas were cryosectioned at 16 microns and mounted on Superfrost Plus slides. For RNAScope labeling, retinal sections were pre-treated according to manufacturer’s instructions for fixed frozen tissue (Advanced Cell Diagnostics, Newark, CA). Probes to *Unc5c* and *Gad2* were from the standard ACD catalog. Probes were hybridized and amplified according to manufacturer’s instructions using the RNAscope Fluorescent Multiplex Kit. Sections were counterstained with DAPI and imaged on a Nikon A1 confocal system. Z-stacks of 0.33 µm steps were acquired using a 60X objective (Plan Apochromat, NA 1.40) with oil immersion. Images were imported into FIJI (Schindelin et al., 2012) and single optical slices were chosen for representative images. For quantification of RNAscope signals, the multi-point tool in FIJI was used to manually mark each nucleus that was positive for the probe. Channels were analyzed individually to blind analysis. Co-expression of markers was used to calculate fraction of *Unc5c* mRNA^+^ cells that expressed *Gad2* mRNA.

### Single cell RNA-seq

Single-cell RNA-seq data from FACS-purified amacrine cells (Yan et al., 2020), adult RGCs (Tran et al., 2019), or developing RGCs (Shekhar et al., 2022) were obtained from the Broad Institute Single Cell Portal as a normalized count matrix, along with the published cluster annotations and UMAP coordinates for each cell. These were loaded into Seurat software (Stuart et al., 2019) for analysis and to generate the gene expression data shown in Supplementary Fig. 2S1. Data from a separate P5 RGC dataset (Rheaume et al., 2018) was accessed via a web-based interface. Cell types within that dataset were identified based on cluster correspondences determined by Shekhar et al. (2022).

To assign amacrine clusters as *Unc5c*-positive or *-*negative (Supplementary Fig. 2S1), we took into account overall expression level across all cells in the cluster (average expression > 0) as well as the fraction of expressing cells within the cluster (a minimum of 20% of cells needed to express the gene to call the cluster positive).

### Immunohistochemistry

For immunohistochemistry, age-matched mouse eyes of both sexes were collected following euthanasia. Eyes were fixed in 4% paraformaldehyde in PBS for 90 min on ice, then corneas and lenses were removed. To label synapses, lenses were removed prior to fixation and eye cups were fixed in 4% paraformaldehyde in PBS for 45 mins on ice. For retinal cross sections, eye cups were cryoprotected in 30% sucrose and 0.02% sodium azide in PBS, frozen in Tissue Freezing Medium (Triangle Biomedical), cryostat sectioned at 20 µm, and placed on Superfrost Plus slides. Sections were blocked in 3% normal donkey serum in PBS with 0.3% Triton X-100 for 1h at room temperature (RT). For retinal flat mounts, retinas were dissected out of eye cups prior to histology, then blocked for 1-2h at RT. Rat anti-mouse Fc block 1:400 (553142, BD Biosciences) was included in the blocking solution for experiments involving mouse antibodies. The following primary antibodies were used: Guinea pig anti-bassoon 1:500 (141004, Synaptic Systems); goat anti-ChAT 1:400 (AB144P, Millipore); rabbit anti-FLRT2 1:25 (ab154023, Abcam); goat anti-FLRT2 1:250 (AF2877 R&D Systems); mouse anti-GAD65 1:1000 (MAB351, EMD Millipore); mouse anti-gephyrin 1:700 (147021, Synaptic Systems); rabbit anti-GFP 1:500 (AB3080P, Millipore); goat anti-GFP 1:500 (ab5450, Abcam); rat anti-Neurofilament-M 1:500 (2H3, Developmental Studies Hybridoma Bank); rabbit anti-Satb1/2 1:500 (2938-1, Epitomics); guinea pig anti-VAChT 1:500 (AB1588, Millipore); mouse anti-Flag 1:1000 (clone M2, F1804, Sigma). Antibodies were diluted in blocking solution, then applied to retinal sections and incubated overnight at 4°C. For flat mounts, 0.02% sodium azide was included in the primary antibody solution and incubation proceeded for 5-7 d at 4°C on a rocker. Sections were washed three times in PBS and fluorophore-conjugated Jackson ImmunoResearch secondary antibodies were applied at 1:1000 for 2h at RT. The following secondaries were used: Alexa Fluor 488 Donkey Anti-Rabbit [711-545-152]; Cy3 Donkey Anti-Goat [705-165-147]; Cy3 Donkey Anti-Guinea Pig [706-165-148]; Alexa Fluor 647 Donkey Anti-Goat [705-605-147]; Alexa Fluor 647 Donkey Anti-Guinea Pig [706-605-148]; Alexa Fluor 647 Donkey Anti-Mouse [715-605-151]; Alexa Fluor 647 Donkey Anti-Rat [712-605-153]. Slides were washed 3 times in PBS then mounted with Fluoromount G. For wholemounts, retinas were washed several times in PBS over 2-4h at RT on a rocker, then secondary antibodies incubated overnight at 4°C on a rocker. Wholemounts were then washed several times on a rocker at RT over 2 hours, mounted on Superfrost slides, coverslipped, and mounted with Fluoromount G.

### Synapse density in IPL

For quantification of synapses, images were acquired using a Nikon A1 confocal system. Z-stacks (106 µm x 106 µm x 0.33 µm step size; 9-15 steps/z-stack) were acquired using a 60X objective (Plan Apochromat, NA 1.49) with oil immersion and 2X optical zoom. Three independent fields of view (FOV) were used for analysis of ON and OFF sublaminae for each animal (n = 3). All FOV were sampled from a similar retinal eccentricity. Images were deconvolved using Nikon Elements Extended Resolution plugin, then imported into ImageJ. Minor brightness/contrast adjustments were made similarly across all images. Maximum intensity projections of three consecutive 0.33 µm z-sections (for a total z-depth of 1 µm) were used for synapse analysis using the Puncta Analyzer plugin for ImageJ (Ippolito and Eroglu, 2010). 3-5 maximum intensity projections were averaged to obtain the number of colocalized puncta per field (dots on graphs). The border between OFF and ON sublaminae was defined based on histologic landmarks (Koh et al., 2018). Puncta Analyzer was used to count synapses based on colocalization of presynaptic and postsynaptic markers as previously described (Ippolito and Eroglu, 2010; Koh et al., 2018). In brief, the projected images were divided into individual channels (presynaptic and postsynaptic markers); background was subtracted for each channel (rolling ball radius = 50); thresholds were adjusted manually to detect individual puncta while minimizing background; and puncta smaller than 4 pixels were filtered out. For all conditions, *N* = 3 animals per genotype, with 3 FOV per animal. Single slices of the z-stacks were chosen for presentation of representative images. Standard error was calculated from 9 FOV from 3 animals per genotype.

### Synapse identification on ooDSGC dendrites

To visualize the distribution of synaptic puncta along dendrites of GFP-labeled ooDSGCs, we utilized a semi-automated method of synapse identification (Della Santina et al., 2013). Images of of synaptic puncta were acquired at a voxel size of 0.104 x 0.104 x 3 µm, deconvolved using Nikon Elements software, and then converted to 8-bit in ImageJ. Using the Simple Neurite Tracer plugin for ImageJ (Longair et al., 2011), we generated a 3D mask of the GFP channel and then used a MATLAB application Object Finder (Della Santina et al., 2013) to identify synaptic puncta within the mask. Object Finder was run with the following settings to segment bassoon and gephyrin puncta: Expected size range = 0.2-5.0 µm; algorithm = iterative thresholding; connectivity = 6; noise estimation = standard deviation; watershed, block search, local noise = true; minimum object intensity = 2x above noise; exclude if single z plane = true. For calling synapses, the minimum overlap between segmented bassoon and gephyrin voxels was set to 50%.

### Analysis of laminar targeting phenotypes

Retinal cross sections at P15-P17 (Figs. 3-4) or P6 and P10 (Fig. 6) were manually inspected for laminar targeting errors using fluorescence microscopy. Confocal z-stacks (0.3-1.0 µm z step) were taken for each region of interest. For each section analyzed, section length was measured in FIJI using the Freehand Line tool. This length was then used to express the number of targeting errors observed in each section as errors per millimeter. Starburst errors were defined as VAChT^+^ arbors ramifying above or below the main S2-S4 VAChT bands. Hb9-ooDSGC errors were defined as GFP^+^ dendrites that were not paired with the starburst scaffold. If the starburst dendrites themselves were targeting an inappropriate layer and GFP^+^ dendrites followed them, this was scored as a starburst error rather than a ooDSGC error.

Satb1 co-labeling (Peng et al., 2017) was used to confirm that the vast majority Hb9-GFP^+^ cells are *bona fide* ooDSGCs at the ages used in this study (Supplementary Fig. 3S2). When Satb1-negative GFP cells were observed, they were always located in the far peripheral retina. Therefore, for Hb9-ooDSGC analysis, the two most peripheral 60X fields of view (FOVs) were not used because these FOVs occasionally include ganglion cells that are not ooDSGCs (Supplementary Fig. 3S2). Phenotype graphs show averaged data from at least 2 sections per animal. Mistargeted arbors were represented graphically by generating fluorescence profile plots through the IPL using FIJI. Raw data were exported to Excel, and for each channel the background signal was subtracted and resulting values were normalized to signal maxima. Images were median filtered (radius = 0.5 pixels) to remove background noise from photomultiplier tube.

### Two-photon guided morphological reconstructions

Isolated whole retinas from adult (∼P40) mice were micro-cut at the dorsal and ventral halves to allow flattening, with dorsal and ventral mounted over a 1–2 mm^2^ hole in nitrocellulose filter paper (Millipore) with the photoreceptor layer side down and stored in oxygenated Ames’ media (maximum 10 h).

Identification, dye loading and imaging of GFP^+^ ooDSGCs was performed as described previously (El-Quessny and Feller, 2021). In brief, GFP^+^ cells were identified using a custom-modified two-photon microscope (Fluoview 300; Olympus America) tuned to 920 nm to minimize activation and bleaching of photoreceptors. The inner limiting membrane above the targeted cell was dissected using a glass electrode. Cell attached voltage clamp recordings were performed with a new glass electrode (4-5 MΩ) filled with internal solution containing the following (in mM): 110 CsMeSO4, 2.8 NaCl, 20 HEPES, 4 EGTA, 5 TEA-Cl, 4 Mg-ATP, 0.3 Na_3_GTP, 10 Na_2_Phosphocreatine, QX-Cl and Alexa-594 Hydrazide dye (pH = 7.2 with CsOH, osmolarity = 290, ECl^-^ = −60 mV). Signals were acquired using Clampex 10.4 recording software and a Multiclamp 700A amplifier (Molecular Devices), sampled at 10 kHz, and low pass filtered at 6 kHz. The Alexa dye freely diffused into the ooDSGC cytoplasm after obtaining a giga ohm (1GΩ) seal and breaking into the cell membrane. Cells were imaged for electrode positioning using two photon excitation at 800 nm. At this wavelength, GFP is not efficiently excited thereby preserving fluorescence for anatomical imaging.

Next, morphological reconstruction and dendritic segmentation were performed as described previously (El Quessny & Feller, 2021). Briefly, 480 x 480 µm Image stacks were acquired at z intervals of 1.0 µm and resampled fifteen times for each stack using a 20X objective (Olympus LUMPlanFl/IR 2x digital zoom, 1.0 NA) and 30kHz resonance scanning mirrors covering the entire dendritic fields of the ooDSGCs. Image stacks of ooDSGCs were then imported to FIJI and a custom macro was used to segment ON and OFF dendrites based on their lamination depth in the inner plexiform layer (ON layer 10-30 µm, OFF layer 35-55 µm depth). Following ON and OFF dendritic segmentation, we used the Simple Neurite Tracer plugin on FIJI to skeletonize and then binarize the ON and OFF dendritic segments for morphological analyses.

### UNC5C gain of function

For AAV production we generated a *ITR*-*CMV-flex-Unc5cHA-WPRE-SV40pA-ITR* construct. A cDNA encoding *Unc5c* (Visser et al., 2015) was tagged at the 3’ end with a 3xHA epitope tag and cloned behind an intronless CMV promoter. Cre-dependence of expression was conferred using the flex-switch strategy (Schnütgen et al., 2003). AAV particles of serotype PHP.Eb were produced from this plasmid by the Duke Viral Vector Core. CAG-flex-tdTomato viral particles (gift of Edward Boyden) was purchased from Addgene (Addgene viral prep #28306-PHPeB).

For AAV administration, heterozygous *Chat^Cre^* P1-2 neonatal mice were anesthetized on ice until unresponsive to toe pinch. Eyelids were cleaned with an ethanol wipe, then a guide hole was punctured at the outer edge of the ora serrata using a 30G needle. A Hamilton syringe was used to inject 0.5-0.75 ul of virus and 1:10 FastGreen was included to visualize injection success. Left eyes were injected with a 5:1 mixture of experimental virus (AAV-flex-Unc5c-HA) and tdTomato reporter virus (AAV-flex-tdTomato); right eyes were injected with the tdTomato reporter virus alone as control. Mice were recovered on a heated pad until they regained pinkness and light aversion. Pups were returned to their home cage with mother after recovery. After 4 weeks, mice were sacrificed and eyes were harvested for histology. Anti-HA staining was performed on each retina to confirm UNC5C protein expression.

### Statistical analysis

All statistical tests were performed using GraphPad Prism software. Non-parametric tests were used in many cases, because large numbers of “0” values (typically in the control condition) led to non-normal data distributions. Sample sizes and P-values are given in figure legends and graphs where applicable; otherwise they are given in Results.

#### Key resources table

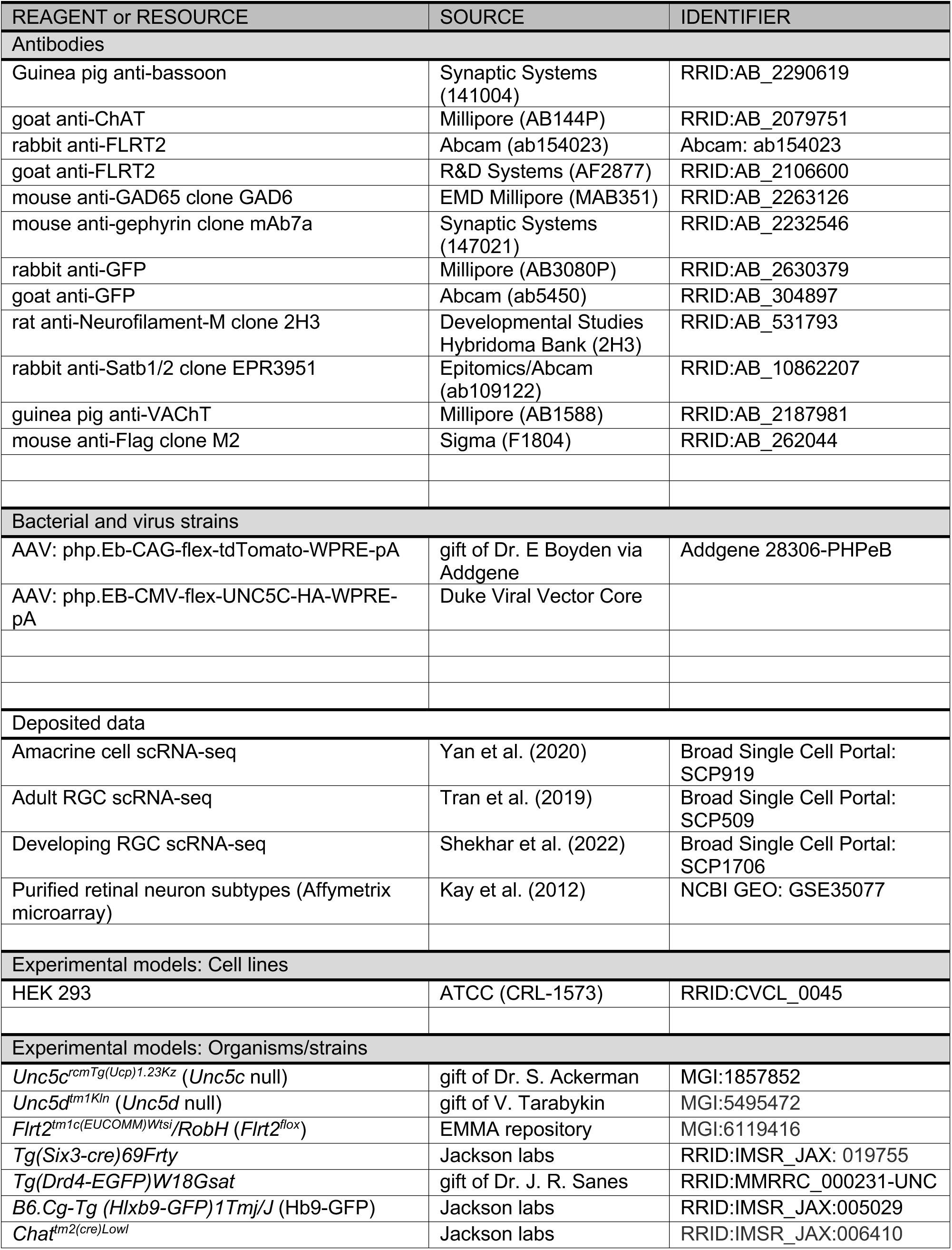
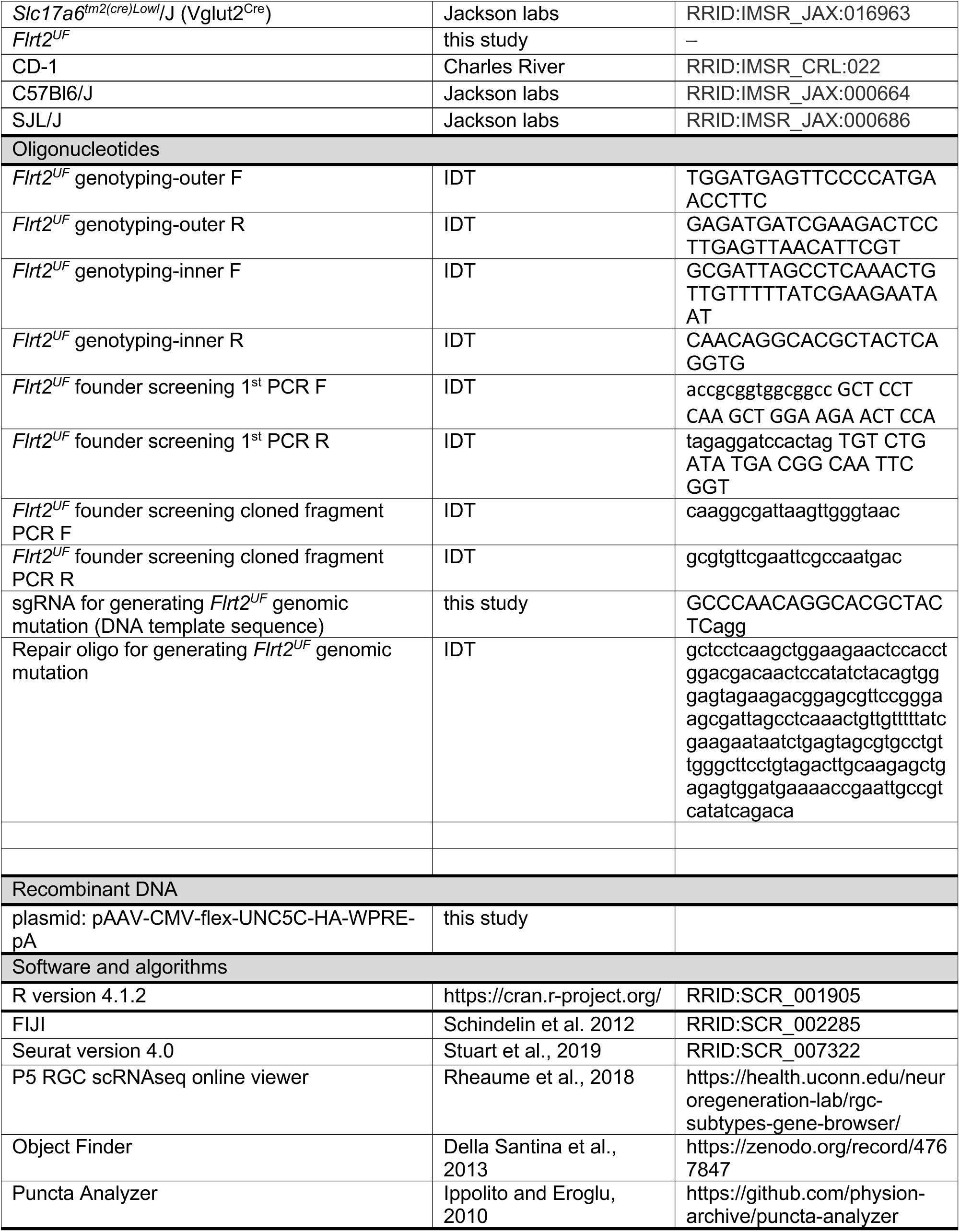

## ACKNOWLEDGEMENTS

This work was supported by the National Eye Institute: EY027998 to C.L.P; EY024694 and EY031445 to J.N.K; EY005722 to Duke University; and EY007551 to University of Houston. Support was also provided by the National Institute of Neurological Disorders and Stroke (NS106756 to M.E.Q.); a Research to Prevent Blindness Unrestricted Grant (to Duke University) and by Glaucoma Research Foundation Shaffer Grant (to L.D.S.). We thank Woj Wojtowicz for reagents, advice, and helpful discussions; Ariane Pendragon and William Kornahrens for technical assistance; Gary Kucera and Cheryl Bock from the Duke Transgenic Mouse Shared resource for their work generating the *Flrt2^UF^*mouse line; Boris Kantor and his team at the Duke Viral Vector Core for assistance with generating the Unc5c-HA construct and AAV virus; and Susan Ackerman, Joshua Sanes, Victor Tarabykin, and David Feldheim for sharing mouse lines.

## Supplementary Figures

**Figure 1S1:**
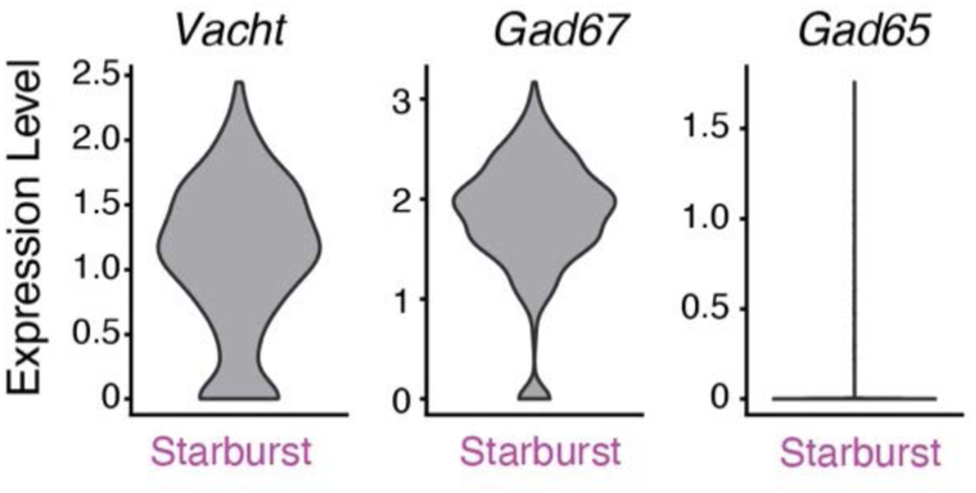
Starburst amacrine cells express GAD67 but not GAD65. Single-cell RNA-seq data from FACS-purified amacrine cells (Yan et al., 2020) was used to assess expression of Gad67 (encoded by the *Gad1* gene) and Gad65 (encoded by the *Gad2* gene) in starburst amacrine cells. Violin plots show expression of Gad67, Gad65, and the starburst marker Vacht (encoded by *Slc18a3* gene) in the cell cluster that was identified in the original publication as corresponding to starburst cells (“AC_17”).

**Figure 2S1:**
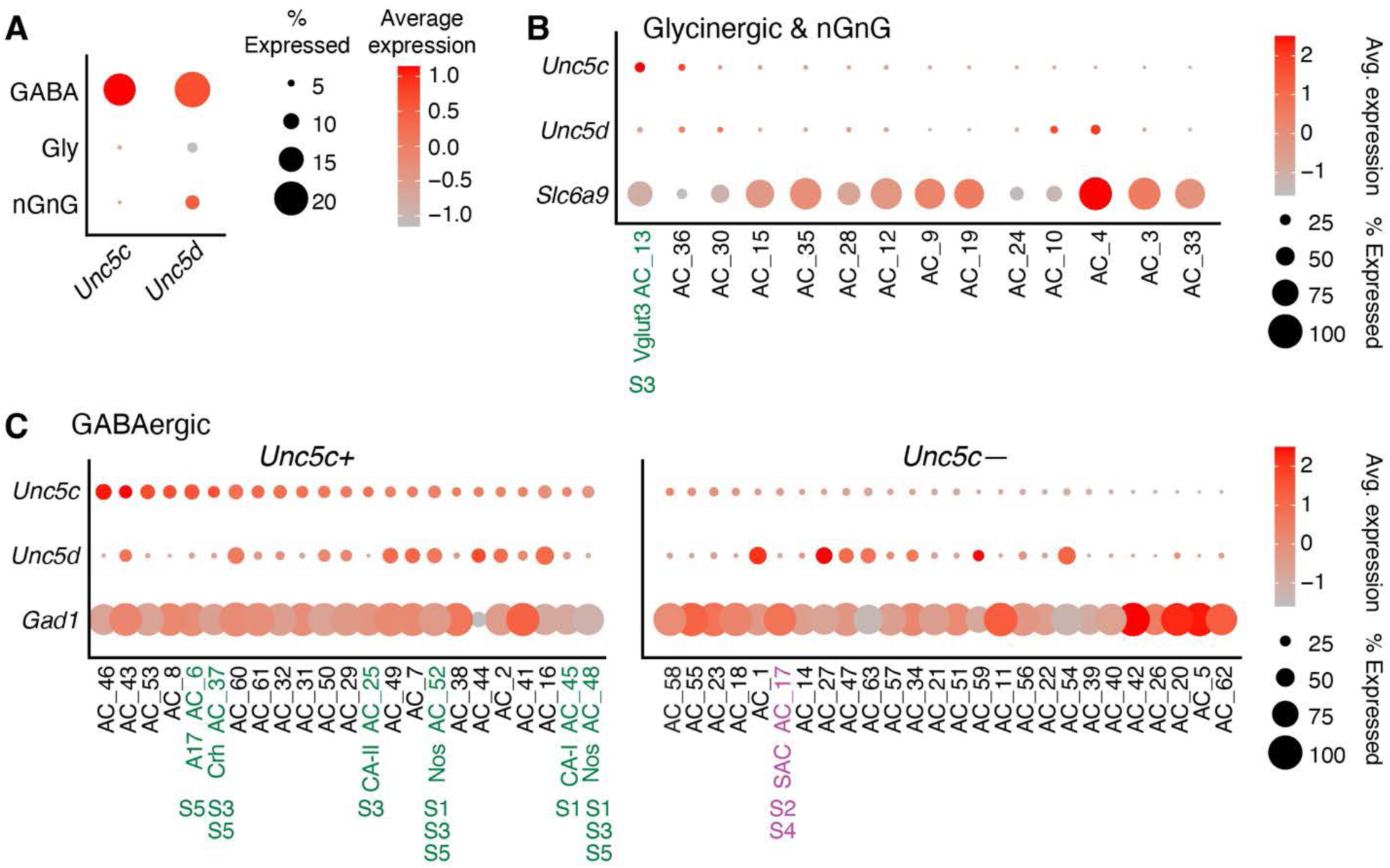
*Unc5c* and *Unc5d* are expressed by GABAergic amacrine cells. Single-cell RNA-seq data from FACS-purified amacrine cells (Yan et al., 2020) was used to assess expression of *Unc5c,* and *Unc5d* genes across amacrine cell types. GABAergic amacrine cells were identified by expression of genes encoding GAD enzymes (*Gad1* and *Gad2*). Glycinergic amacrine cells were identified by expression of *Slc6a9*, encoding the glycine transporter GLYT1. Amacrine types that are neither GABAergic nor glycinergic (nGnG) were identified by absence of all 3 marker genes. Each cluster from the published dataset was assigned to one of these three classes. **A**. Dotplot showing expression of *Unc5c* and *Unc5d* by the broad amacrine cell classes. *Unc5c* and *Unc5d* are predominantly expressed by GABAergic amacrine cells. **B,C.** Expression of *Unc5c* and *Unc5d* by individual amacrine cell clusters. B, glycinergic and nGnG clusters; C, GABAergic clusters. Green and magenta text marks clusters corresponding to known cell types, with known arborizations in particular IPL layers as indicated. Clusters were assigned as *Unc5c*-positive (C, left) or *Unc5c-*negative (C, right) based on average gene expression across the cluster as well as the fraction of expressing cells (see Methods).

**Figure 3S1:**
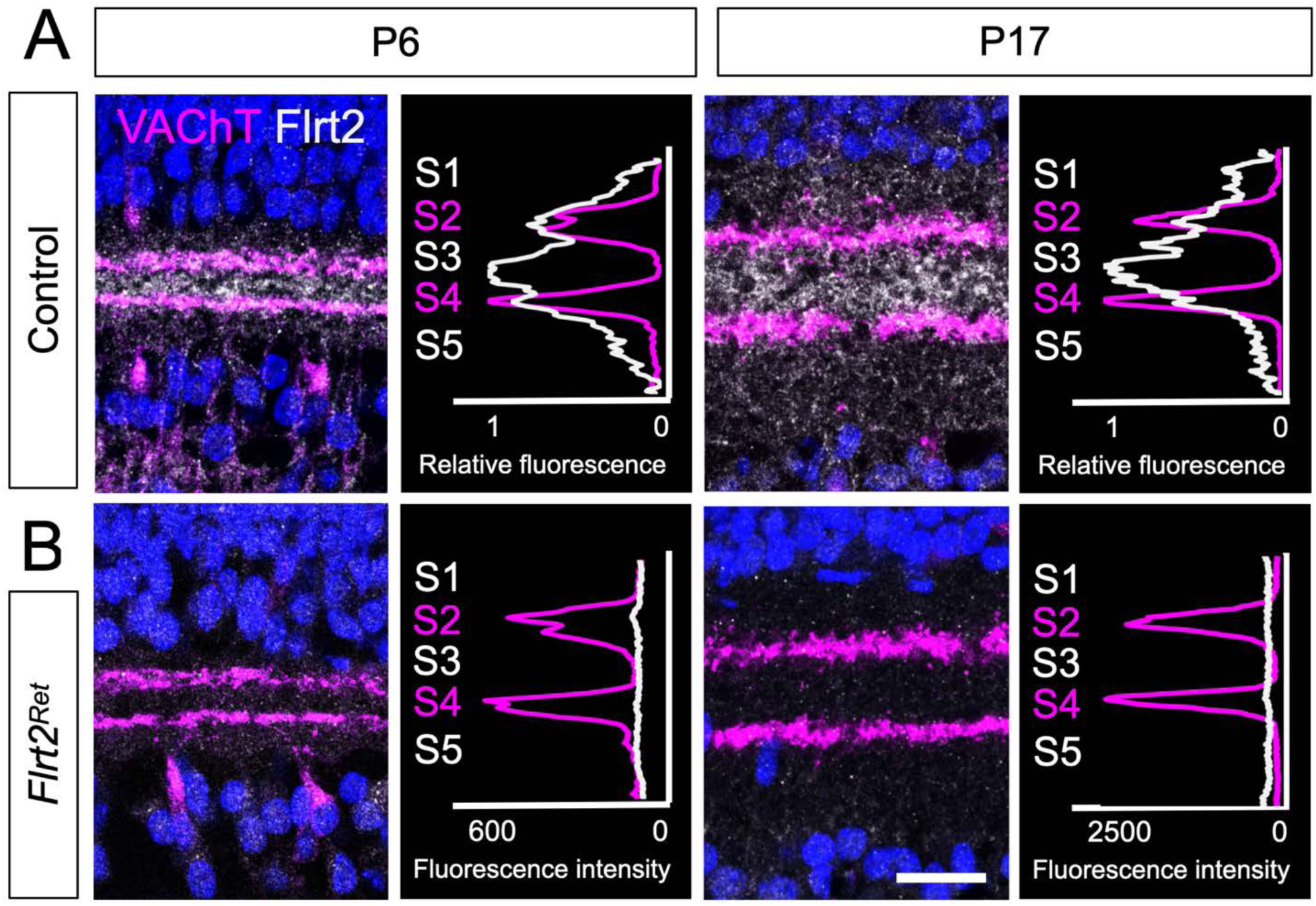
FLRT2 protein is absent in *Flrt2^Ret^* mutants. **A**. FLRT2 protein expression in retinal cross sections stained with anti-VAChT for starburst arbors (magenta) and anti-FLRT2 (white). At P6 (left panels), wild-type FLRT2 protein is expressed primarily in the DS circuit layers in S2 and S4, with an additional peak in S3. A similar expression pattern is seen at P17 (right panels). This expression is consistent with previous reports (Visser et al., 2015). **B**. FLRT2 protein is absent in *Flrt2^Ret^* mutants at both P6 (left panels) and P17 (right panels). Due to the lack of staining in *Flrt2^Ret^*, absolute fluorescence intensity is plotted for *Flrt2^Ret^* panels. Scale bar, 20 µm.

**Figure 3S2:**
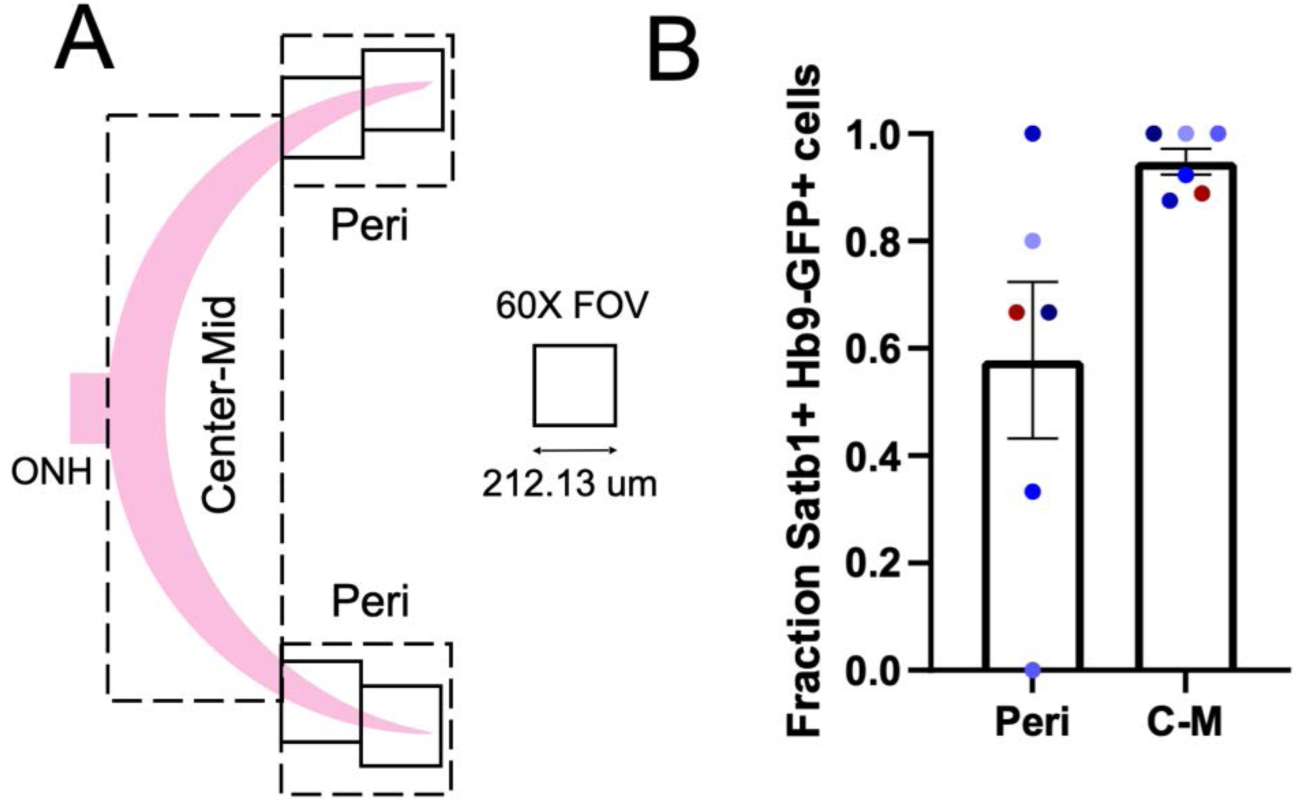
Hb9-GFP reliably labels ooDSGCs. **A**. Schematic of sampling area used for phenotype and colocalization analysis. Retinal cross-sections were analyzed under 60X magnification. Field of view (FOV) size for each image was 212.13 µm x 212.13 µm. The first two FOVs were designated as the peripheral (Peri) regions, with the region to either side of the optic nerve head (ONH) out to the periphery designated as the Center-Mid region. **B**. Retinal cross-sections were stained with anti-GFP and anti-Satb1/2, a marker of ooDSGCs (Peng et al., 2017). The fraction of GFP^+^ cells expressing the ooDSGC marker was determined separately for Center-Mid (C-M) and Peri regions. In Hb9-GFP retinas, 57.8% of GFP^+^ cells in the periphery were co-labeled by Satb1, indicating 42.2% of GFP^+^ cells in this region are not ooDSGCs. Therefore, the periphery of Hb9-GFP retinas was excluded from analysis of laminar targeting phenotypes. GFP^+^ cells in the Center-Mid retina were reliably ooDSGCs (94.8%), so all Center-Mid GFP^+^ cells were included in analysis. N = 6 animals. Blue points, P15, red points, P10. Datapoints from individual animals share same color tone in Peri and C-M graphs.

**Figure 3S3:**
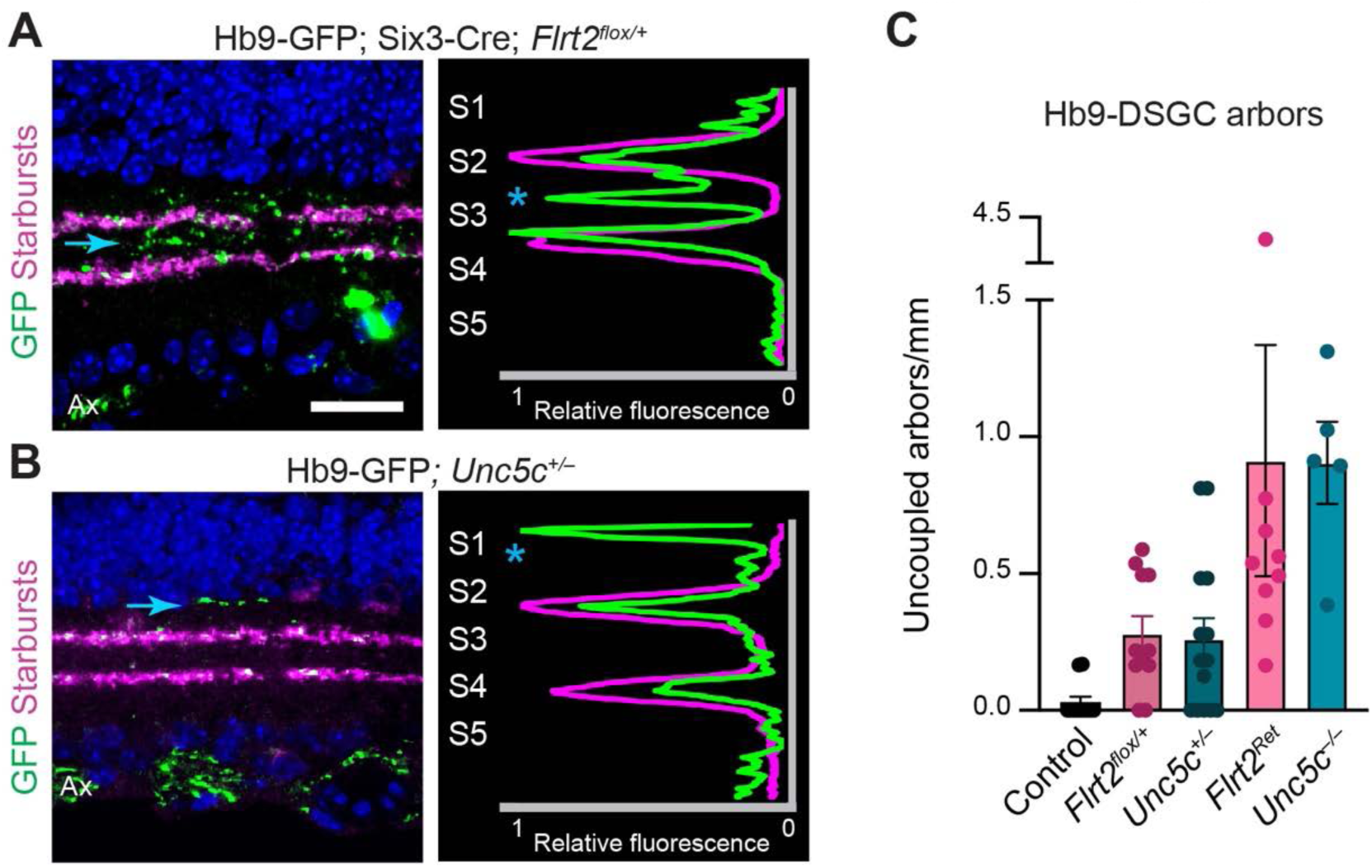
ooDSGC dendrite targeting phenotypes in *Flrt2* and *Unc5c* heterozygotes. **A-B**. Representative retinal cross-sections from P15 Hb9-GFP mice of the specified *Flrt2* or *Unc5c* genotype, stained with anti-GFP to reveal ooDSGC arbors and anti-VAChT to label starburst arbors. Both *Flrt2* heterozygotes (A) and *Unc5c* heterozygotes (B) show laminar targeting errors (arrows) resembling those observed in homozygous mutants (main Fig. 3). Ax, axons in nerve fiber layer. Scale bar, 20 µm. **C**. Summary of Hb9-ooDSGC errors in Six3-Cre; *Flrt2^flox/+^* heterozygotes and *Unc5c^+/–^* heterozygotes. Homozygous mutant data is replotted from Fig. 3E for comparison. Heterozygous mice had more errors than controls (p = 0.008), but fewer than homozygous mutants (p = 0.006). Statistics: Kruskal-Wallace test (p = 4 x 10^-7^) followed by Dunn’s post-hoc test. Due to similar magnitude of heterozygous and homozygous error rates, genotypes were pooled for analysis.

**Figure 3S4:**
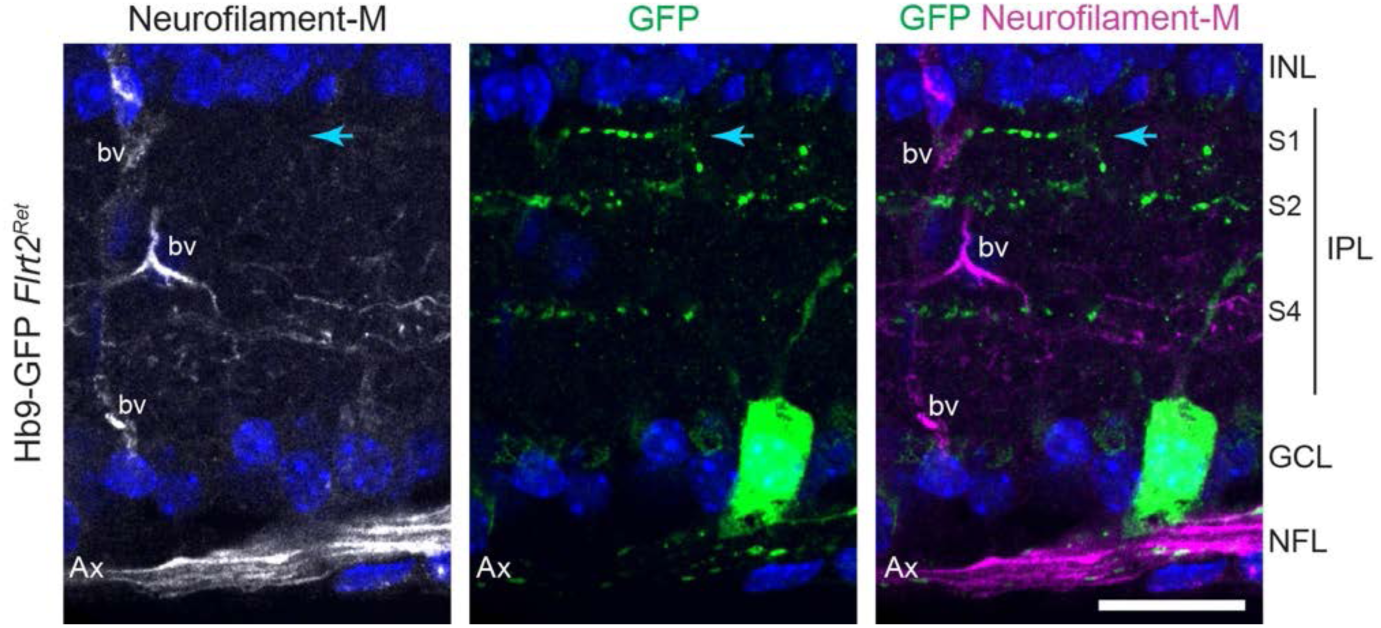
ooDSGC arbors projecting to ectopic IPL regions are not axons. Representative image of retinal cross section from P15 *Flrt2^Ret^* mutant, co-stained for GFP to label Hb9-DSGCs and neurofilament-M to label axons. Antibodies to neurofilament-M label retinal ganglion cell axons (Ax) within the nerve fiber layer (NFL). bv, blood vessels labeled nonspecifically by anti-mouse secondary antibody. An ectopic Hb9-ooDSGC arbor (blue arrow) is not labeled by antibodies to neurofilament-M. Scale bar, 20 µm.

**Figure 3S5:**
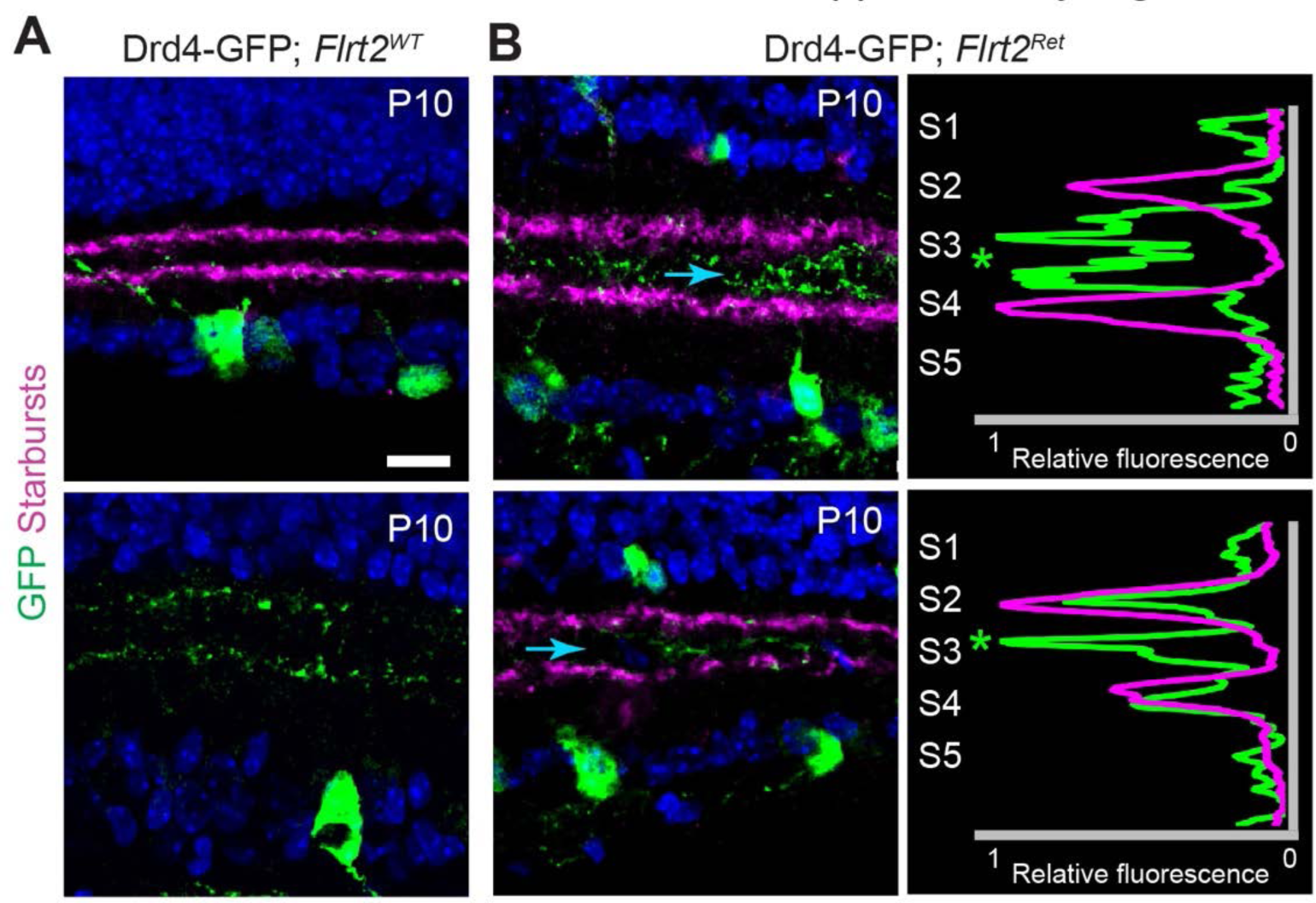
Laminar targeting errors in Drd4-GFP; *Flrt2^Ret^* mutants. **A**. Two representative cross-sectional images (top, bottom) of P10 Drd4-GFP retinas from *Flrt2^WT^* controls. GFP^+^ ooDSGC dendrites cofasciculate with starbursts (magenta) in S2 and S4. **B.** Two examples (top, bottom) of Drd4-ooDSGC laminar targeting errors in *Flrt2^Ret^* mutants. We were unable to obtain larger sample sizes due to genetic linkage between the GFP transgene and the *Flrt2* locus. Ectopic arbors (blue arrow in left panel and asterisk in right panel) projected into S3 in a diffuse (top) or stratified (bottom) manner. Note that P10 was analyzed due to downregulation of GFP at later timepoints, such that we could not reliably identify ooDSGCs past P10. Scale bar, 20 µm.

**Figure 3S6:**
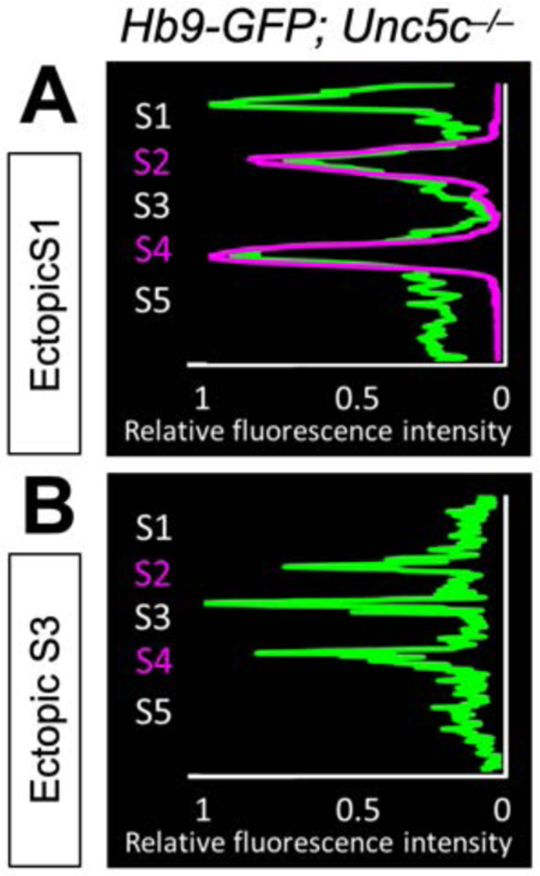
*Unc5c^−/−^* error profile plots. **A-B**. Representative profile plots from Hb9-GFP *Unc5c^−/−^*retinas through ectopic projections in S1 (*A*, top) and S3 (*B*, bottom). Green = GFP; Magenta = starbursts.

**Figure 4S1:**
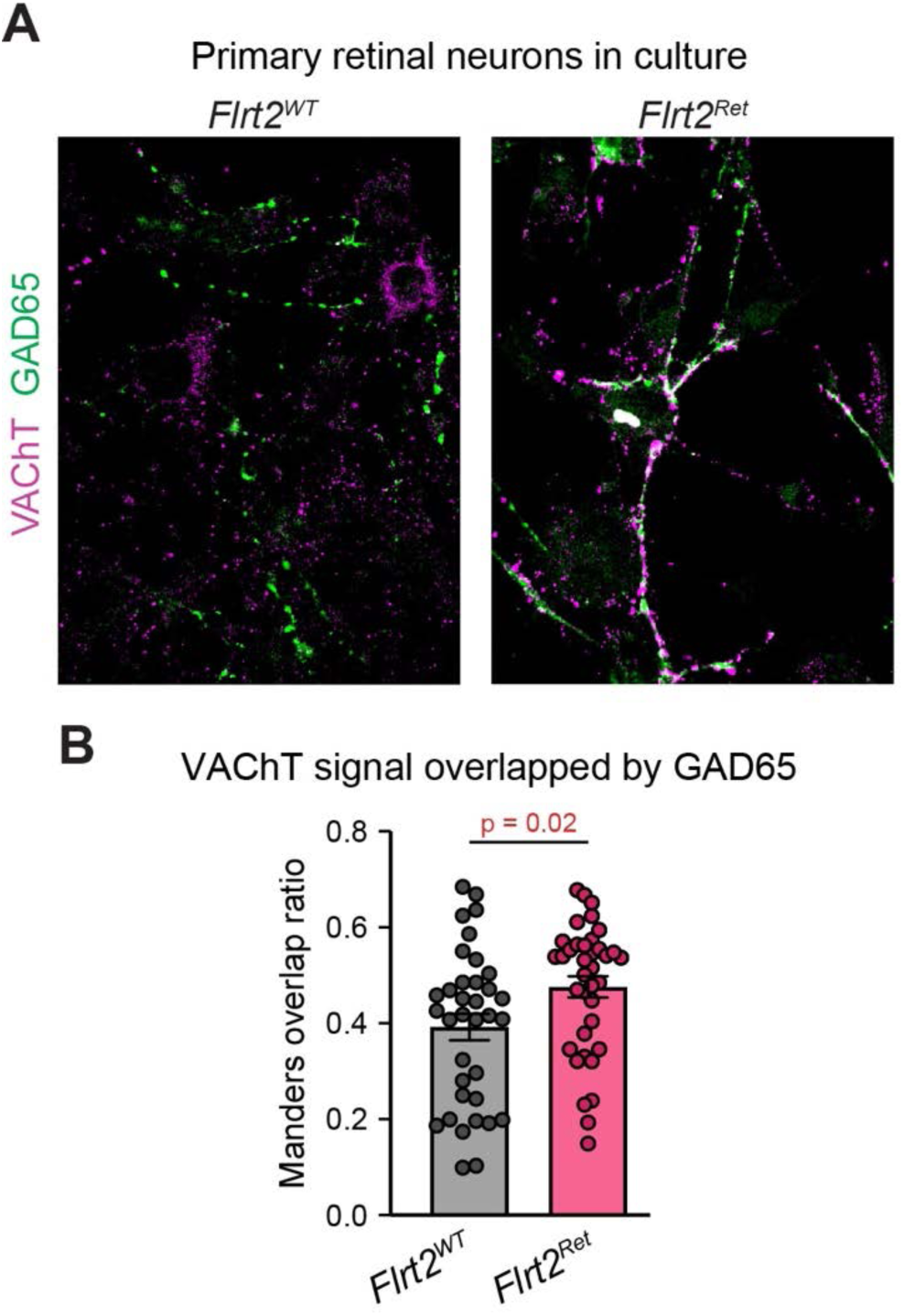
Loss of *Flrt2* increases overlap of starburst and GAD65^+^ amacrine arbors. **A**. Primary retinal neuron cultures were prepared from wild-type (left) or *Flrt2^Ret^* (right) mutant mice at P1-2, and fixed 1 week later. Starburst arbors were labeled by anti-VAChT (magenta); non-starburst GABA neurons were labeled by anti-GAD65 (green). Sites where starburst and GAD65^+^ arbors overlap appear white in color. **B.** Quantification of VAChT^+^ territory colocalized with GAD65^+^ signal using the Manders M1-M2 ratio methodology (Manders et al., 1992). Graph shows ratio of VAChT pixel intensities in pixels containing GAD65 signals vs. all pixels. Pixel values were normalized prior to calculating the Manders ratio (see Methods). Sample sizes: n = 35 (wild-type) or 37 (*Flrt2^Ret^*) images from two separate culture experiments. Dots, measurements from individual images. Error bar, S.E.M. Statistics: two-tailed unpaired T-test.

**Figure 4S2:**
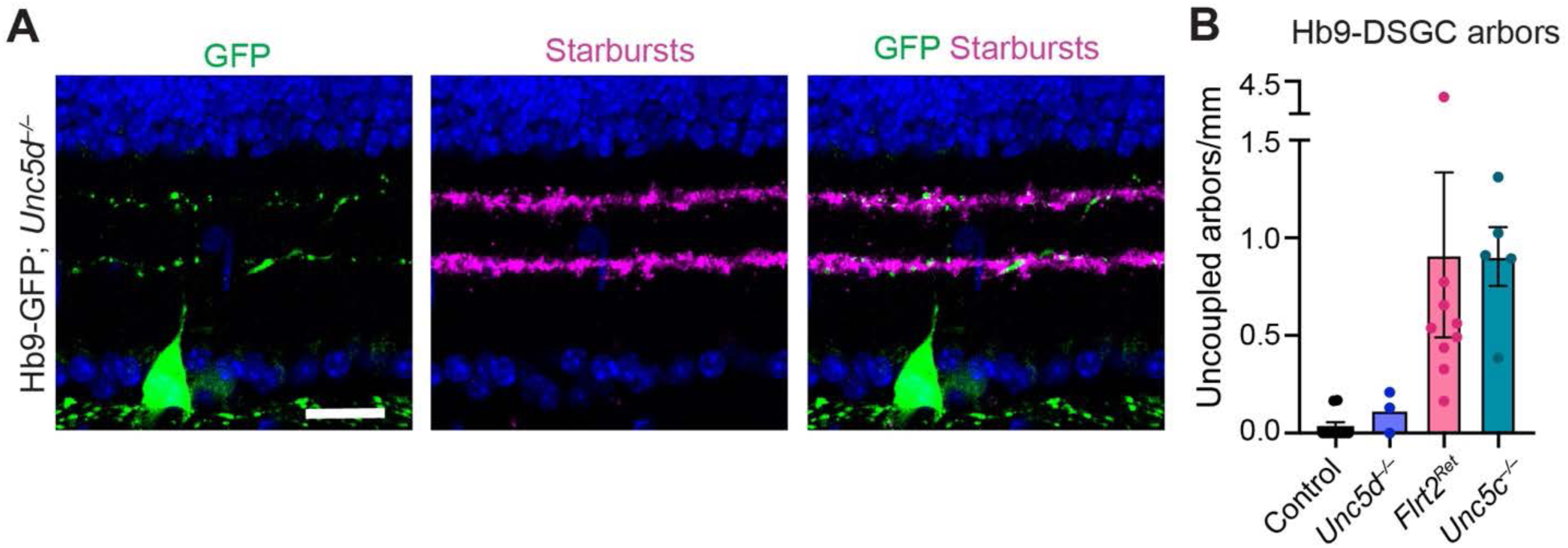
Laminar targeting of ooDSGCs is unchanged in *Unc5d* mutants. **A**. Representative cross-sectional image from P15 *Unc5d^−/−^*mutant. Green, Hb9-ooDSGCs; magenta, starburst dendrites labeled by anti-VAChT. *Unc5d^−/−^* retinas exhibit normal lamination in S2 and S4 (green, left panel), where they associated with starburst scaffold (magenta, center panel; merged image, right panel). Scale bar = 20 um. **B.** Frequency of uncoupled Hb9-ooDSGCs in *Unc5d* mutants (n = 3) was similar to controls, suggesting *Unc5d* is not involved in Hb9-ooDSGC laminar targeting. For comparison *Flrt2^Ret^* and *Unc5c* mutants are replotted from Fig. 3.

**Figure 5S1:**
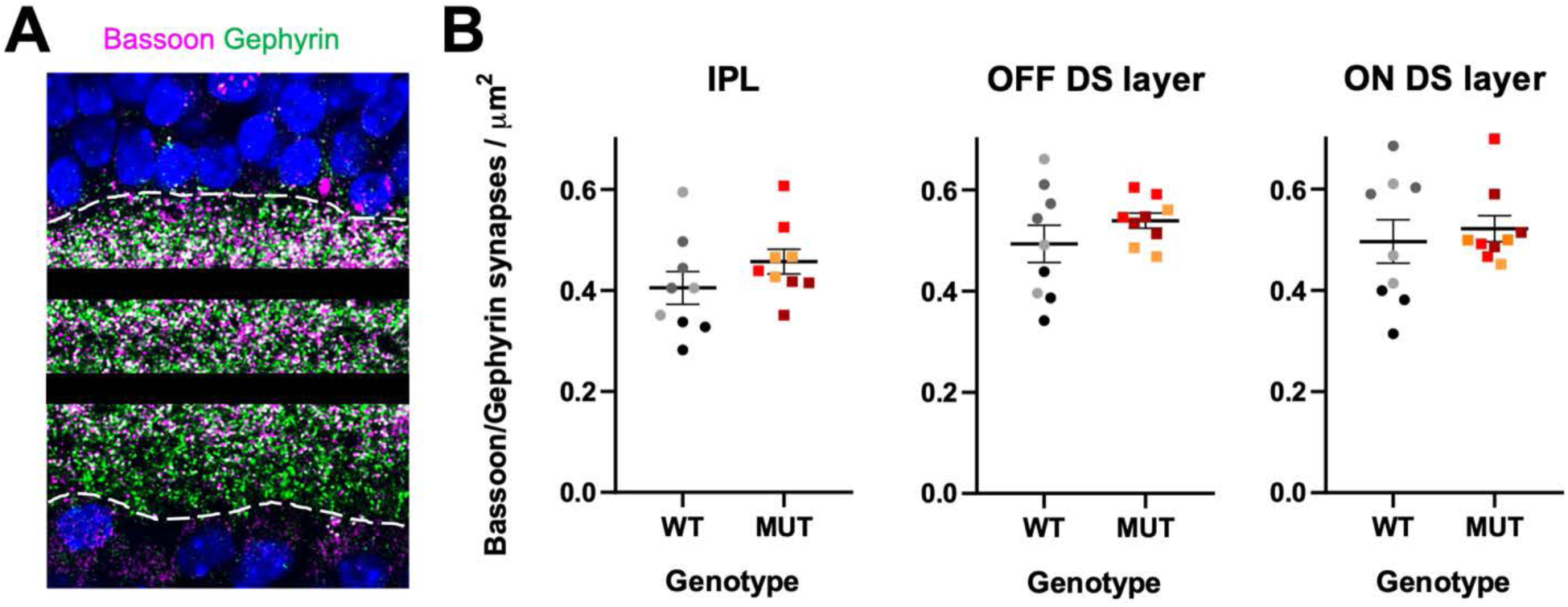
Synapse density across IPL. **A**. Representative image of inhibitory synapse labeling in a P17 retinal cross-section. Pre-synaptic sites labeled by anti-bassoon (magenta), post-synaptic sites labeled by anti-gephyrin (green); colocalization (white) indicates putative synapses. Synapses were counted throughout the IPL (within region contained by broken white line) or in masks of the OFF DS layer (top black bar) or ON DS layer (bottom black bar). **B**. Quantification of synapse density in wild-type and *Flrt2^Ret^* mutants from images similar to A. Synapse density was unchanged across the entire IPL in mutants (left panel). There was also no difference in synapse density in either the OFF DS layer (center panel) or ON DS layer (right panel). Data from P17 mice. N = 3 animals per genotype; 3 FOVs analyzed per animal (indicated by dots of same color); each FOV was produced by averaging 3 Z-slices covering 1 µm of Z distance.

**Figure 6S1:**
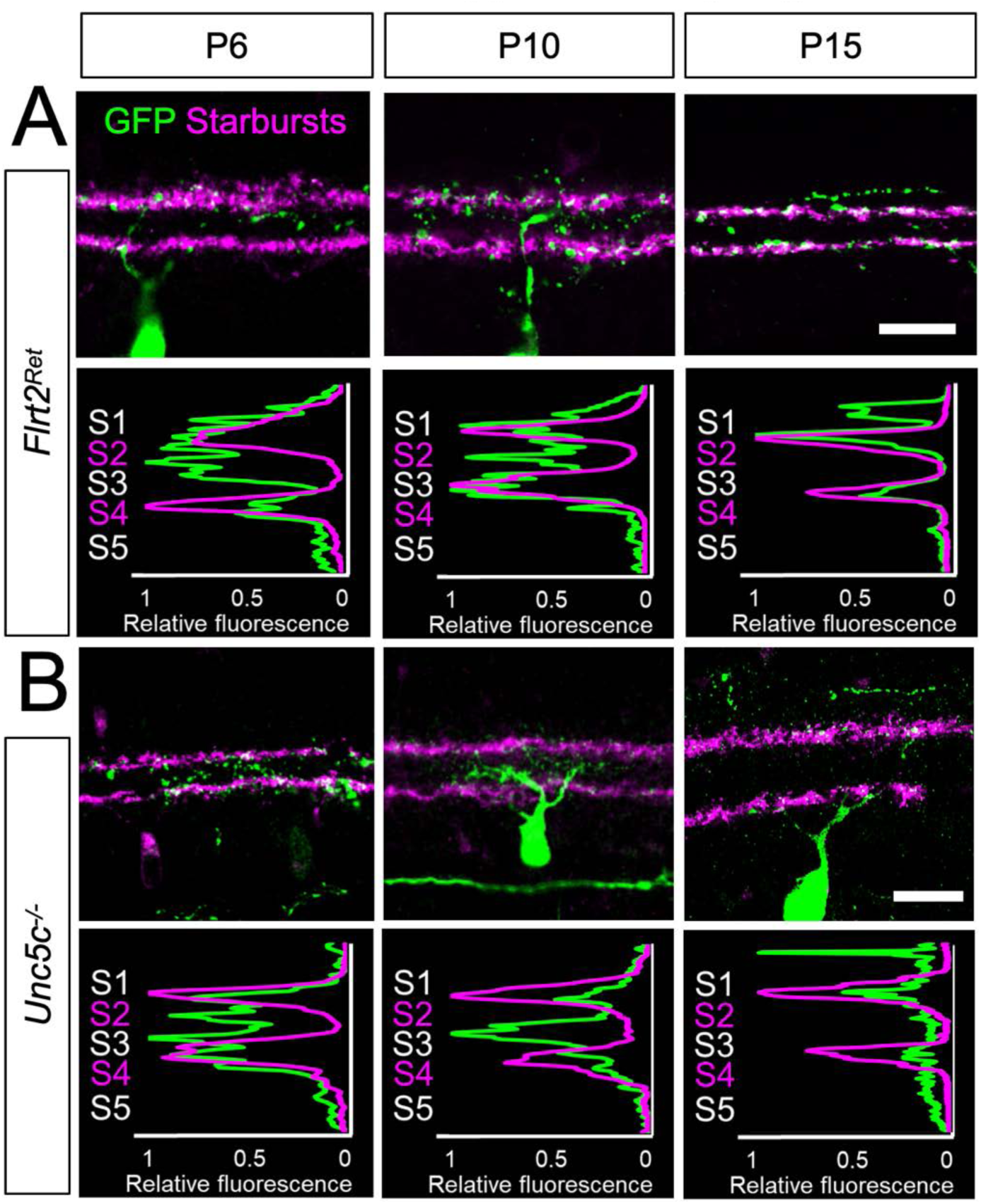
*Flrt2^Ret^* and *Unc5c^−/−^* laminar targeting at P6, P10, and P15. Retinal cross-sections from *Flrt2^Ret^* (A) and *Unc5c^−/−^* (B) mutants at indicated ages, showing examples of ooDSGC dendrites that were scored as mistargeted (i.e. uncoupled from the starburst scaffold). Images illustrate the phenotypes quantified in Fig. 6E. Mutants were littermates of the wild-type animals shown in Fig. 6A-D. Green, anti-GFP; magenta, anti-VAChT. Scale bar = 20 um.

**Figure 7S1:**
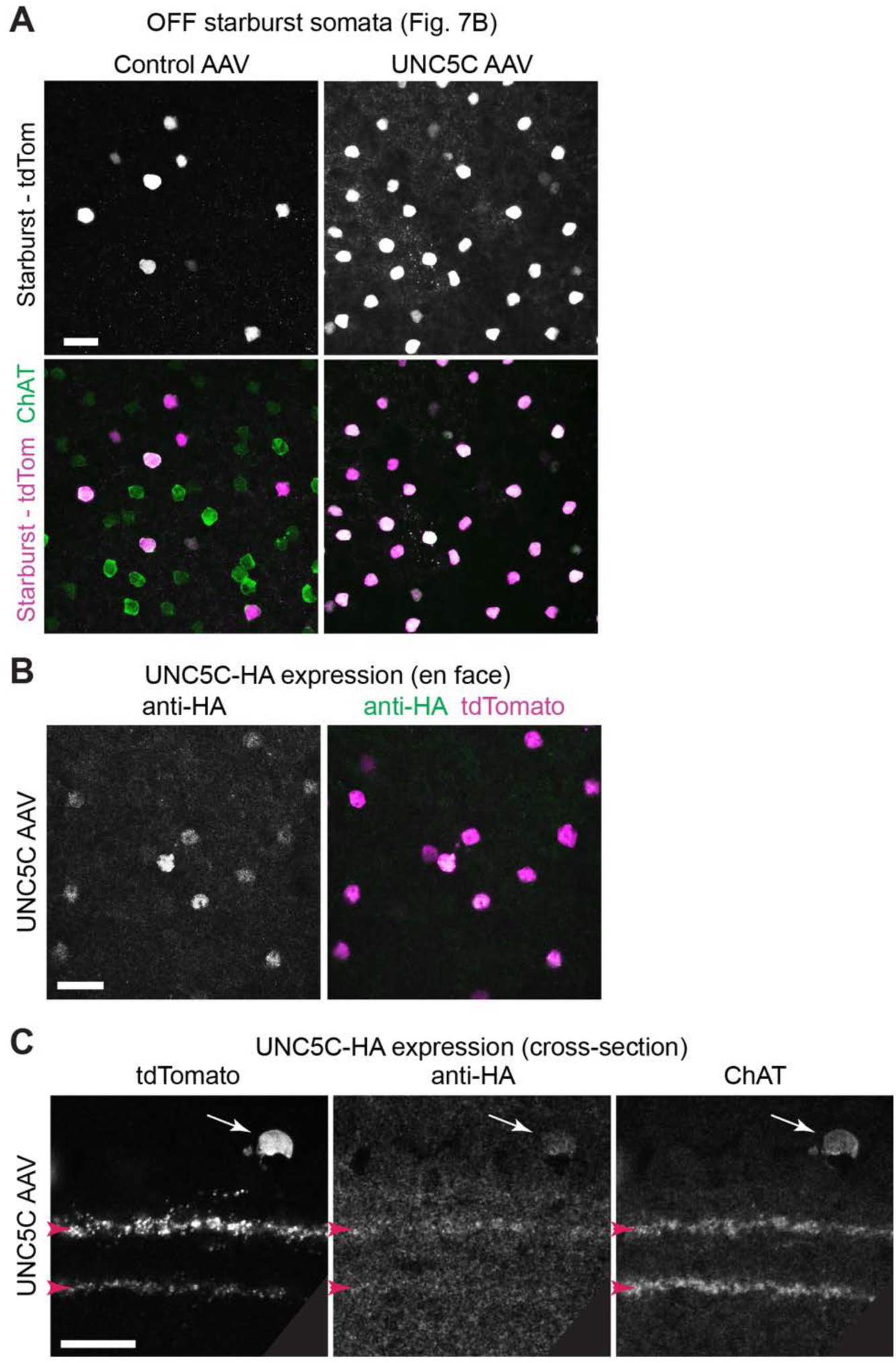
AAV-mediated expression of UNC5C-HA in starburst cells. **A**. En-face views of AAV-infected retinal wholemounts, at the level of the INL, showing somata of transduced OFF starburst cells. In both control and Chat-UNC5C AAV conditions, all transduced cells (labeled by anti-tdTomato) are Chat^+^ starburst cells, supporting the specificity of starburst labeling. Images are from the same retinal wholemounts depicted in Fig. 7B, but a wider view to give a sense of how many starburst cells were infected in these retinal regions. These images show that the difference in tdTomato^+^ starburst IPL labeling between control and UNC5C AAV retinas (Fig. 7B) cannot be explained by a difference in the number of transduced starburst cells. Indeed, the UNC5C-expressing region had a far higher fraction of tdTomato^+^ starbursts, but a notably lower density of starburst arbors in the IPL. **B,C**. Validation of UNC5C protein expression by starburst cells transduced using the Chat-UNC5C AAV strategy. B, En-face view of wholemount retina at INL level, showing that the vast majority of tdTomato^+^ starburst cells also express UNC5C-HA. C, cross-sections showing that UNC5C-HA localizes to ChAT^+^ IPL sublayers (red arrowheads) in infected retinal regions. Arrow, soma of infected ChAT^+^ OFF starburst cell co-expressing UNC5C-HA and tdTomato.

